# Pervasive homeobox gene function in the male-specific nervous system of *Caenorhabditis elegans*

**DOI:** 10.1101/2025.05.13.653874

**Authors:** Robert W. Fernandez, Angelo J. Digirolamo, Giulio Valperga, G. Robert Aguilar, Laura Molina-García, Rinn M. Kersh, Chen Wang, Karinna Pe, Yasmin H. Ramadan, Curtis Loer, Arantza Barrios, Oliver Hobert

## Abstract

We explore here how neuronal cell type diversity is genetically delineated in the context of the large, but poorly studied male-specific nervous system of the nematode *Caenorhabditis elegans.* Mostly during postembryonic development, the *C. elegans* male adds 93 male-specific neurons, falling into 25 cardinal classes, to the predominantly embryonically generated, sex-shared nervous system, comprised of 294 neurons (116 cardinal classes). Using engineered reporter alleles, we investigate here the expression pattern of 40 phylogenetically conserved homeodomain proteins within the male-specific nervous system of *C. elegans,* demonstrating that in aggregate, the expression of these homeodomain proteins covers each individual male-specific neuron. We show that the male-specific nervous system can be subdivided along the anterior/posterior axis in HOX cluster expression domains. The extent of our expression analysis predicts that each individual neuron class is likely defined by unique combinations of homeodomain proteins. Using a collection of newly available molecular markers, we undertake a mutant analysis of five of these genes (*unc-30, unc-42, lim-6, lin-11, ttx-1)* and identified defects in cell fate specification and/or male copulatory defects in each of these mutant strains. Our analysis expands our understanding of the importance of homeobox genes in nervous system development and function.

## INTRODUCTION

The generation of molecular maps of animal brains has tremendously advanced over the past few years, with whole brain atlases now existing for several bilaterian model system species (e.g.[1–3]). Molecular brain maps raise a host of questions: Can the multidimensional complexity of individual neuron types be reduced to simpler molecular descriptors? Are there common themes in the mechanisms that generate the enormous diversity of cell types which define each animal nervous system? A tentative answer to both questions has recently emerged in the brain of the nematode *Caenorhabditis elegans*: first, the analysis of expression of the entire family of homeodomain transcription factors (encoded by a total of 102 homeobox genes) has shown that each of the 118 distinct neuron classes of the nervous system of the hermaphrodite can be described by unique combinatorial codes of homeodomain expression [4, 5]; and, second, mutant analyses of homeobox genes over the past few decades have revealed that homeobox genes indeed regulate the acquisition of specific neuronal identities – not just in *C. elegans*, but in many other animal species as well (reviewed in [6]).

To further investigate how extensively homeobox genes define distinct neuronal cell types, we turned to the little explored nervous system of the *C. elegans* male. Based on sex-specific patterns of blast cell proliferation, sex-specific execution of cell death programs and sex-specific transdifferentiation, male animals generate an additional set of 93 neurons compared to the hermaphrodite [7–11]. Owing to lineage history, overall morphology and synaptic connectivity, these 93 neurons can be subdivided into 25 “cardinal classes” [7–11](**Table 1**). Two cardinal classes are located in the head (CEM and MCM neuron classes), two are located in the ventral nerve cord (CA and CP), and all others are located in several distinct ganglia in the tail of the animal, where they form a closely intertwined set of circuits that control various aspects of the complex male copulatory behavior [12–14]. While many of the 25 cardinal classes are composed only of either unilateral neurons or bilateral neuron pairs, four cardinal classes – the CA and CP ventral cord neurons, and the tail ray sensory classes RnA and RnB - are subdividable into a multitude of different subclasses that are clearly distinguishable by connectivity and molecular markers [15–17].

**Table 1:**
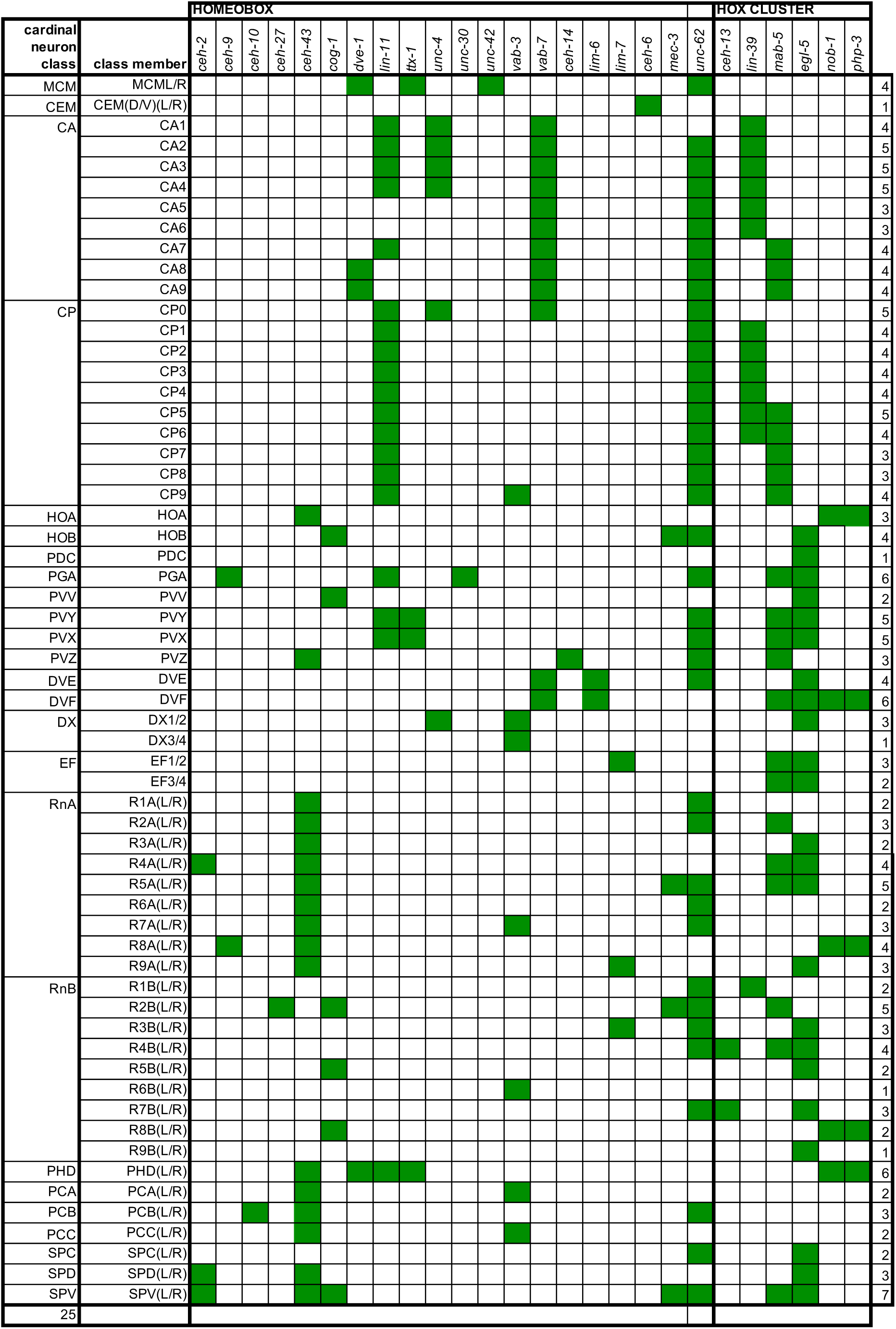
Summary of expression of the homeobox genes across the male *C. elegans* nervous system. This table summarizes the imaging data from **Fig 1**. Sites of expression were identified by crossing homeobox reporter alleles into the NeuroPAL (*otIs669* or *otIs696*) landmark strain. Panneuronally expressed *ceh-44* and *ceh-48* are not shown here. Numbers on the right show how many homeodomain proteins are expressed in a given neuron class.

In contrast to the mostly embryonically generated, sex-shared nervous system of *C. elegans*, where neurons are generated after a series of rapid cell divisions, most male-specific neurons are generated from quiescent neuroblasts, generated initially as specialized epithelial cells in the embryo. In early larval stages, these neuroblasts lose their specialized epithelial features, re-enter the cell cycle and generate the vast majority of male-specific neurons, as well as other male-specific cell types [7, 18]. These distinctive patterning mechanisms could trigger terminal differentiation programs that are distinct from those that generate sex-shared neurons during embryogenesis. In the most extreme version of such scenario, these differentiation program may even rely less on the homeobox gene family that is so important in patterning the embryonic, sex-shared nervous system.

Only some very limited hints towards the involvement of homeobox genes in male-specific neurons exist. Male-specific CEM neurons were previously described to require the *unc-86/Brn3* POU homeobox gene for their proper differentiation [19–22]. In the ventral cord, HOX cluster genes *lin-39* and *mab-5* control the differentiation of two classes of male-specific motor and interneurons, the CA and the CP neurons, while the HOX cluster gene *egl-5* controls the proper differentiation of a subset of ray sensory neurons [23–27]. Beyond these hints of homeobox gene function in certain male-specific neurons, very little is known about the expression or function of homeobox genes in the many remaining male-specific neurons. We have set out here to address this gap in knowledge by making use of a large toolbox of strains that express *gfp-*tagged homeobox genes, a resource that we previously used to investigate homeodomain protein expression throughout the entire nervous system of the hermaphrodite [4, 5].

Cellular sites of gene expression patterns in the male-specific nervous system have been notoriously difficult to identify based on the occasionally variable position of neuronal cell bodies in the male tail and the absence of reporter landmarks. This problem was recently overcome by the introduction of NeuroPAL, a multicolor fluorescent transgene in which each individual male-specific neuron class can be reliably identified through non-GFP-based fluorescent landmarks, which can be overlaid with a GFP-tagged reporter strain [28]. NeuroPAL color maps have been established for both the hermaphrodite and the male nervous system [28, 29].

The paucity of well-described fluorescent reporter-based marker genes has hampered the study of neuronal differentiation programs for which such markers are often of critical importance. Compounding this lack of markers is also the current absence of a complete scRNA-seq atlas of the male-specific nervous system. However, the recent systematic mapping of neurotransmitter identities in the male-specific nervous system has begun to mitigate this problem [15]. In addition, our recent reporter-based analysis of neuropeptide-encoding genes in the hermaphrodite nervous system [30] has provided additional tools, in the form of neuropeptide reporter alleles, to identity more cell fate markers in the male-specific nervous system. We exploit these tools here to generate a collection of cell fate markers of male-specific neuron identities and to assess the impact of homeobox gene function on the differentiation programs of several male-specific neuron types. The analysis conducted here in this paper leads us to conclude that despite the distinctive patterning mechanisms of male-specific neurons, homeobox genes play a role in male-specific neuron differentiation that appears to be as predominant as in the sex-shared nervous system.

## MATERIAL AND METHODS

### Strains

A list of strains used in this study is provided in **S2 Table**.

### *C*. *elegans* genome-engineered strains

Many reporter alleles generated in this study were engineered using the CRISPR/Cas9 system, by SunyBiotech, using an SL2::GFP::H2B reporter cassette (indicated by the *syb* allele name in **S2 Table**). Some of these strains have been recently described [4, 5, 30, 31], other have been generated specifically for this study (**S2 Table**).

All gene deletion alleles were designed to remove the entire coding region of the gene, and include the following:

A null allele for *unc-42*, *ot1187*, was generated injecting the following crRNAs and repair template to completely remove the coding region of *unc-42*:

crRNA (GCTCATtgtgtgagtgaaag), crRNA (tctcactgatagactaatgt), ssODN (ATCCCTTCAGAGCCATACTTCTCACTACTACCACCATCATAGAATCAAGACCTGAAATCGACCTAAAAAA).

Due to issue with their mating efficiency, multiple molecularly identical null alleles of *lin-11* and *lim-*6 have been generated by injecting separate strains carrying reporter alleles for terminal identity markers, using the following crRNAs and repair templates:

*lin-11(ot1026, ot1497, ot1444, ot1521, ot1472, ot1483)*, crRNAs (attgagaagggagtaaaagg and CGTGGAATACTCCTGTATGT), ssODN

(TTCGTGGTCGttcttcttcttcttctcctcctcctTACAGGAGTATTCCACGTTCGTGTAGTTTTTCTTC)

*lim-6 (ot1699, ot1700, ot1701, ot1702)*, crRNAs (TGTGTTTTGTAGAAGACCGG and GAAAAGCAAAATAAAGCGGG), ssODN

(gctcctgctctctctctctgtgttttgtagaagacgctttattttgcttttcacctcatattatttattt)

### Neuronal identification using NeuroPAL

Sites of reporter gene expression were determined using the NeuroPAL landmark strain (*otIs669* and *otIs696* alleles) and male tail atlases previously described [28, 29]. The identity of each neuronal type was identified by comparing the color, size, and location of each neuron relative to one another. A detailed protocol to neuronal identification using NeuroPAL can be found at: https://www.hobertlab.org/neuropal/

### Microscopy and mutant analysis

To prepare animals for imaging, a small agarose pad (5%) was cast on a standard imaging slide and worms were mounted and immobilized using a solution of 100 mM of sodium azide (NaN3). Images were acquired either using confocal laser scanning microscopes (Zeiss LSM880 and LSM980) or wide-field microscopy (Axio Imager Z2). Images were processed and analysed using the Zen (Zeiss) or Fiji [32] imaging software. All reporter reagents and mutants were imaged at 40x using fosmid or CRISPR reagents, unless otherwise specified.

For determining the expression pattern of homeobox genes or terminal identity markers, representative maximum intensity projections are displayed in grey scale, with gamma and histogram adjustments for visibility. For mutant functional analysis, representative maximum intensity projections are shown in inverted grey scale. In case of all-or-nothing changes, a qualitative analysis using three qualifiers was used to score mutants; ON - the signal was still present in the mutant, OFF – the signal was lost, DIM – the signal was still present but drastically reduced. When changes were not all-or-nothing, a qualitative approach was used and the signal intensity was extracted using either Zen or Fiji.

To assay homeobox gene expression pattern at different larval and adult stages, animals were picked from a plate containing a mixed population of different stages. Different larval stages were differentiated according to size and well-known anatomical markers. Furthermore, animals were sexed under a high magnification microscope and only male larvae were included in the analysis.

The NeuroPAL images provided in supplementary figures are pseudo-colored according to [28, 29].

### Serotonin antibody staining

Anti-serotonin immunofluorescence was performed as previously described [33]. Briefly, worms were fixed overnight (ON) in 1.5 ml microfuge tubes at 4C in 4% paraformaldehyde in PBS; rinsed 3x in PBSTx (0.5% Triton X-100 / PBS), then incubated ON at 37C with gentle mixing in 5% beta-mercaptoethanol in TrisTx (1% TX-100 / 0.1 M Tris, pH 7.4). Rinsed twice in TrisTx, then once in collagenase buffer (CB: 1mM CaCl_2_ / TrisTx), then digested with 2000 Units/ml Collagenase type IV (Sigma C5138) in CB until a few adult worms fragmented (typically 30-45 min). Rinsed three times PBSTx, incubated in 1% BSA in PBSTx for 1-2 hrs at room temp (RT). Then incubated ON at RT in 1:100 anti-serotonin (Rabbit antiserum, Sigma S5545) in 1% BSA/ PBSTx. Rinsed three to four times in 0.1% BSA/ PBSTx over 1-2 hr at RT. Then incubated ON at RT (in the dark) in 1:100 secondary antibody (Goat anti-Rabbit IgG, TRITC-conjugated). Rinsed 3-4x in 0.1% BSA/ PBSTx over 1-2 hr at RT. Viewed and photographed with an Olympus BX60 upright fluorescence microscope equipped with a Magnafire CCD camera.

### Mating assays

Mating assays were performed on 9 cm NGM plates in which a bacterial lawn of 15 uL of OP50 was placed in the middle containing 30 *unc-51(e369)* hermaphrodites. Males were tested at one day of adulthood with 1-day-old *unc-51(e369)* hermaphrodites picked the night before as L4s. Each male was tested for 15 minutes. During this time, all steps of mating were scored in one or more hermaphrodites. Assays were replicated at least twice on different days and with different sets of males. Videos of mating events were recorded at 2fps using LoopBio and visualized with QMPlay 2 to analyze the following steps of mating:

*Response:* A male was scored as responding to mate contact if it placed its tail ventral side down on the hermaphrodite’s body and initiated the mating sequence by backing along the hermaphrodite’s body to make a turn. The response efficiency was calculated by dividing 1 (response) by the total number of contacts made with the mate before responding.

As a more sensitive measure of the quality of response, we scored hesitation during response. Hesitation is a switch in direction between forward and backward locomotion from the time the male establishes contact with the mate to the first turn (or to location of vulva if this occurs without the need of a turn).

*Scanning:* A single scan was scored as the journey around the hermaphrodite’s body away from and returning to the vulva position. The first scan was counted as the journey from the point of first contact to the hermaphrodite vulva position. A scan was considered continuous if locomotion was maintained in the backward direction without switching direction or pausing (regardless of pause duration).

*Turning:* Measured as proportion of good turns (number of good turns divided by total number of turns performed by the male) until location of vulva. A turn was considered good if it happened continuously while the worm was scanning backwards the tip of the hermaphrodite body to continue scanning the other side of the hermaphrodite without losing contact, switching direction or pausing before the turn.

*Location of vulva (LOV):* A male was considered successful in locating the vulva when they stop scanning at the vulva position to try to insert the spicules. The LOV efficiency was calculated by dividing 1 (LOV) by the total number of times the male passes by the vulva without stopping there.

*Molina maneuvers:* A continuous single maneuver was scored as the journey away from the vulva in forward locomotion, to a distance bigger than two tail-tip lengths, and return to the vulva in backward locomotion. Any visible pause during forward or backward locomotion was considered a STOP regardless of its duration. The category of discontinuous maneuver ‘switching’ was scored as a change in direction of locomotion while travelling away or towards the vulva without reaching it.

*Tail contact loss:* Number of contact loss was scored as previously described [34]; i.e. the number of times that a male lost tail contact with the hermaphrodite during the mating trial (without counting male detachment after ejaculation).

## RESULTS

### Homeodomain protein expression analysis

The *C. elegans* genome codes for 102 homeobox genes, 80 of them conserved throughout the animal kingdom (**S1 Table**). Since no complete scRNA-seq dataset exists so far for the male-specific nervous system of *C. elegans*, we examined homeobox gene expression using a reporter gene approach, making use of a resource of *gfp-*tagged homeobox gene loci whose expression we previously analyzed in the context of the hermaphrodite [4, 5]. This approach has the advantage over scRNA-seq analysis that the direct fusion of *gfp* to the respective homeobox gene visualizes protein, rather than mRNA expression, thereby capturing posttranscriptional (e.g. translational) gene regulatory events.

We analyzed the expression of half (40) of the 80 conserved homeodomain proteins, covering all main homeodomain subclasses (Antp-like, Prd-like, POU, LIM, SIX *etc.*). We assessed expression of all male-specific neurons in the head, ventral nerve cord and tail (**Fig 1**; **Table 1, S1 Table**). For the identification of the neuronal sites of expression, we used the NeuroPAL transgene which differentially labels all neuron classes in the male-specific nervous system [29](**S1 Fig**).

**Fig 1:**
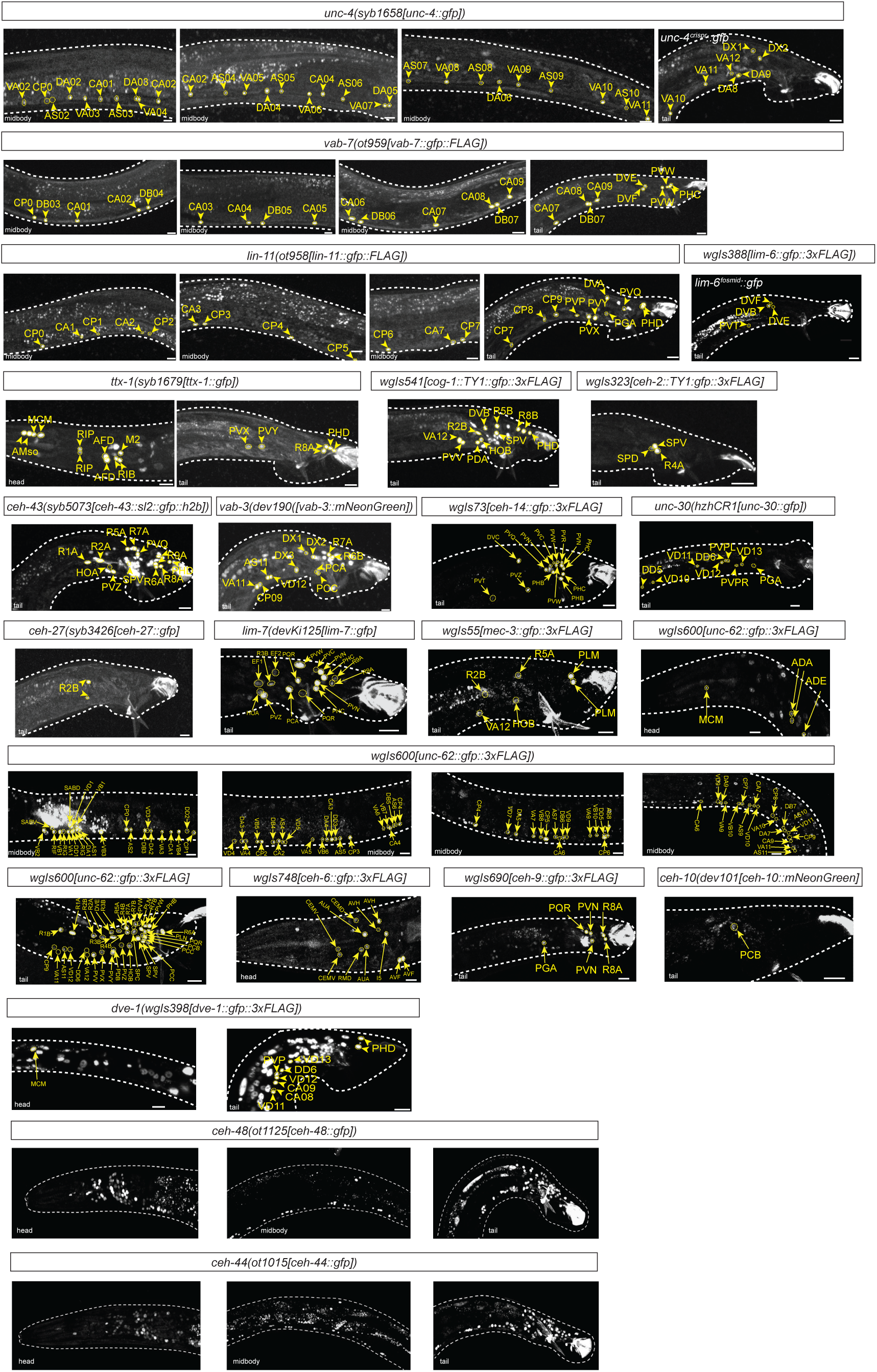
Representative images of homeobox reporter alleles. Representative images of homeobox reporters examined in this paper, which are either fosmid-based or CRISPR/Cas9-engineered alleles. Neuronal sites of expression were identified through overlap with NeuroPAL [28, 29]. The corresponding images showing overlap with NeuroPAL colors are in **S1 Fig,** with the exception of the pan-neuronal *ceh-44* and *ceh-48*. Expression patterns are summarized in **Table 1**.

For 12 of the examined 40 homeodomain proteins, we observed no expression in male-specific neurons (summarized in **S1 Table**). As expected from their pan-neuronal expression in the hermaphrodite [35], two genes, the CUT homeobox genes *ceh-44* and *ceh-48* are panneuronally expressed (**Fig 1**). All other 26 homeodomain proteins show restricted expression in male-specific neurons (**Fig 1**, **Table 1**). We found that each male-specific neuron class expresses at least one homeodomain protein, therefore matching the complete coverage of all neuron classes in the hermaphrodite by homeobox genes. Most homeobox genes are expressed in small subsets of male-specific neuron classes, sometimes exclusively in single neuron classes (*unc-42/Prop1* in MCM, *unc-30/Pitx* in PGA). Exceptions to very restricted homeodomain proteins patterns are the six HOX cluster proteins, as well as their frequent co-factor UNC-62/Meis, each of which are broadly, yet still highly cell type-specifically expressed throughout the male-specific nervous system. Other than the two of the AbdB-type HOX cluster genes, *php-3* and *nob-1* [36], which recently duplicated specifically in *Caenorhabditiae,* no two homeobox genes show the exact same expression pattern (**Fig 1**, **Table 1**).

Each individual male-specific neuron co-expresses, on average, three homeobox genes (range: 1 to 7, excluding the panneuronal CUT homeobox genes). There appear to be no preferential partnership of any two homeodomain proteins, with the exception of the expected, frequent association of UNC-62/Meis expression with a HOX cluster gene. This is expected because of the evolutionary ancient biochemical partnership of HOX and Meis proteins [37]. However, there are also clear cases where either HOX or Meis are expressed independently of one another, as also observed in other organisms [38]. We describe HOX protein expression patterns in more detail in a separate section below.

Strikingly, of the cardinal 25 male-specific neuron classes, there are only two sets of neuron classes that cannot be distinguished by unique combinations of the homeobox genes we examined (even though they can be distinguished by neuropeptides, **Table 2**): The lineally related PCA and PCC share the same homeobox code (*ceh-43* and *vab-3)* as do the lineally related PVX and PVY (*lin-11* and *ttx-1,* plus 3 HOX cluster genes)(**Table 1**). As reflected by their shared name (and their shared lineage), the PCA and PCC neuron class, as well as the PVX and PVY neuron classes are also anatomically similar to one another. Nevertheless, since we only analyzed half of all conserved *C. elegans* homeobox genes, we anticipate that a complete examination may attach unique codes to all neuron classes.

**Table 2:**
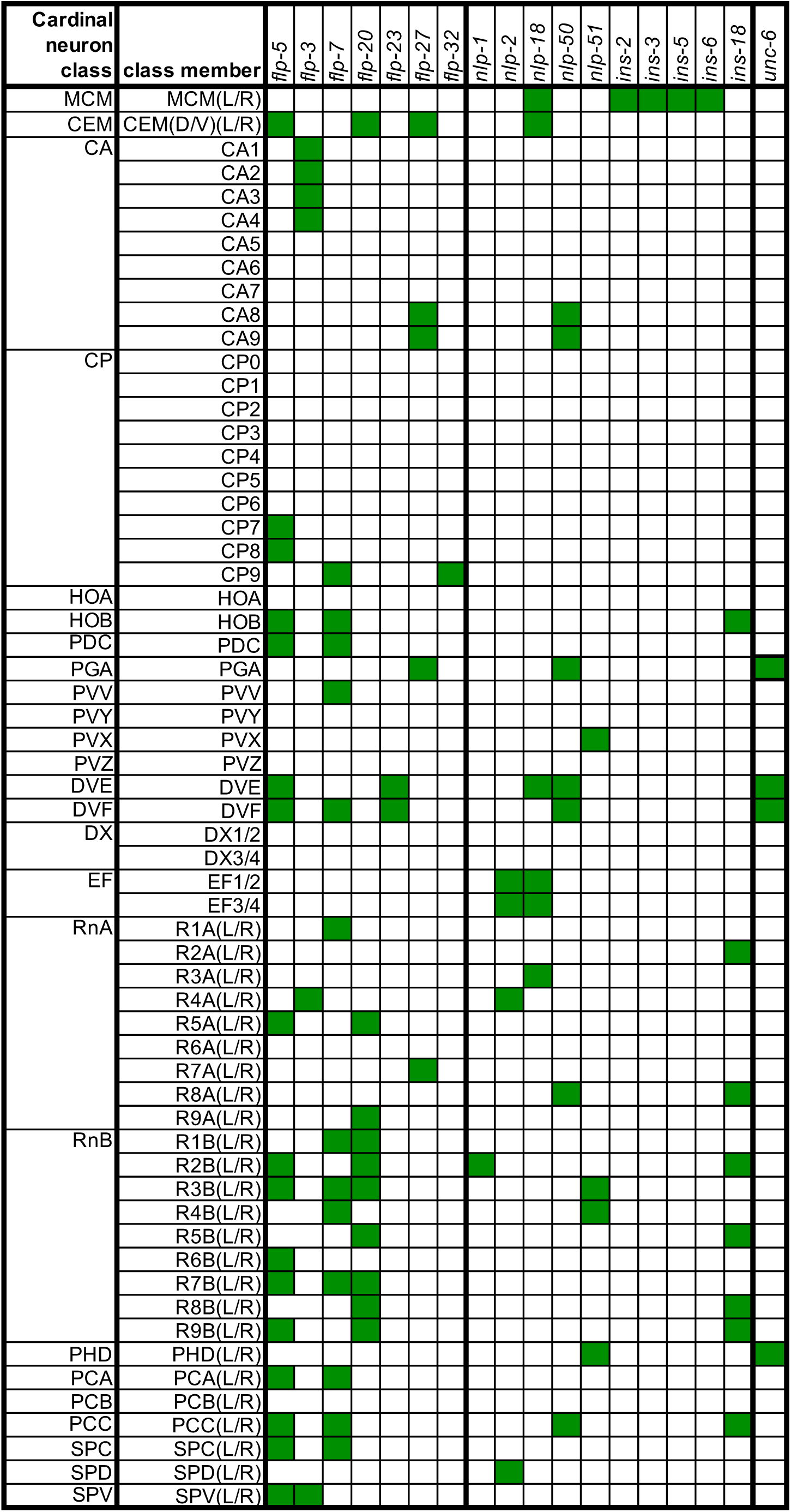
Summary of molecular markers for male-specific neurons. This table summarizes the imaging data from **Fig 4**. Sites of expression were identified by crossing homeobox reporter alleles into the NeuroPAL (*otIs669* or *otIs696*) landmark strain.

Within each cardinal class that can be clearly subdivided into subtypes (RnA, RnB, CAn and CPn neurons), we also find homeobox codes for several, but not all molecularly previously defined subtypes (**Fig 1**, **Table 1**). Specifically, within the ventral nerve cord, we observe that homeobox genes subdivide male-specific CA and CP neuron classes into subclasses, in accordance with earlier studies using cell fate markers [23] and recent neurotransmitter mapping studies [15]. Similarly, in the male tail, the A- and B-type ray neuron classes, each composed of 9 class members, can be subdivided into various subclasses based on homeobox gene expression (**Fig 1**, **Table 1**), again in accordance with earlier studies using other markers [15, 16].

We discovered novel subtypes with the DX and EF neuron class, each of which is composed of up to 4 class members (*lim-7/Islet:* EF1/2, not EF3/4; *unc-4:* DX1/2, not DX3/4). Conversely, neuron classes with previously much appreciated subclass diversity are “unified” by the expression of particular homeobox genes: all CA neurons express the Eve/Evx-type homeobox gene *vab-7,* all CP neurons express the *lin-11* LIM homeobox genes and all A-type ray neurons express Dlx-type homeobox gene *ceh-43* (**Fig 1**, **Table 1**).

### HOX cluster genes show anteriorly/posteriorly patterned expression in the male-specific nervous system

One set of homeobox genes that warrant a separate consideration are the *C. elegans* HOX cluster genes (**Fig 2A and 2B**). In the context of animal nervous systems, the analysis of HOX gene expression and function has largely focused on the ventral nerve cord of invertebrates and spinal cord of vertebrates where HOX cluster genes are differentially expressed along the anterior/posterior axis, showing a remarkable match to their chromosomal localization (“co-linearity”)(reviewed in [39–42]). In the *C. elegans* hermaphrodite, HOX gene expression extends posteriorly beyond the ventral nerve cord into various tail ganglia, which express the AbdB-homologs *egl-5, php-3* and *nob-1* (reviewed in [39]). In the male, the expression of a subset of HOX cluster had already been examined in some restricted regions of the male tail (e.g. [43]; reviewed in [39]), but no comprehensive and comparative analysis was yet available. Using reporter alleles for all HOX cluster genes, we note the following themes of HOX genes in the adult male nervous system (**Fig 2B and S2; Table 1**):

1. The vast majority of mature, male-specific neurons in the ventral cord and tail ganglia express at least one HOX cluster gene. This is in striking contrast to neurons in the anterior head ganglia, which express very few HOX cluster genes (**Fig 2B**).
2. Along the male ventral nerve cord, the postembryonically added, male-specific CA and CP neurons show similar patterns as the sex-shared neurons: Concordant with their chromosomal location, *lin-39/Scr* and *mab-5/Antp* show an ordered expression with *lin-39/Scr* being expressed in more anterior CA and CP class members, and *mab-5/Antp* in more posterior CA and CP class members.
3. In male all tail ganglia, which contain a manifold increase in neuron number of all different types (sensory, inter-and motor neurons) compared to hermaphrodites, domains of HOX gene expression show a notable co-linearity with their genomic arrangement: there is no expression of the anterior HOX cluster gene *lin-39/Scr*, while *mab-5/Antp* expression is observed in more anterior parts of the ganglia but then peters out. *egl-5/AbdB* expression predominates, with the most posterior *AbdB* paralogs *nob-1* and *php-3* being more enriched in the most posterior neurons (**Fig 2B**; **Table 1**).
4. Colinearity of HOX gene expression is recapitulated, albeit imperfectly, in the “microcosm” of the ray neurons, which innervate the sensory ray structure of the worm that are aligned in an anterior-to-posterior manner along the male bursa (inset in **Fig 2A**).
5. The *ceh-13/Lab* homeobox gene represent a curious case in the *C. elegans* HOX cluster. *ceh-13* is the *C. elegans* homolog of the most anterior HOX cluster gene of other metazoans, Labial in flies and *HoxA/B/C/D1* in vertebrates, but it has switched its chromosomal order with *lin-39,* the *C. elegans* Dfd/Scr homolog (**Fig 2A and 2B**)[36, 44]. Nevertheless, in the embryo, *ceh-13* expression pattern is indeed enriched in the anterior part [45, 46]. However, unlike with the other HOX cluster genes, an anterior/posterior gradient is not evident in the context of the hermaphrodite ventral nerve cord [4]. We rather find that in males, *ceh-13/Lab* is expressed in several ray neurons in the tail.

**Fig 2:**
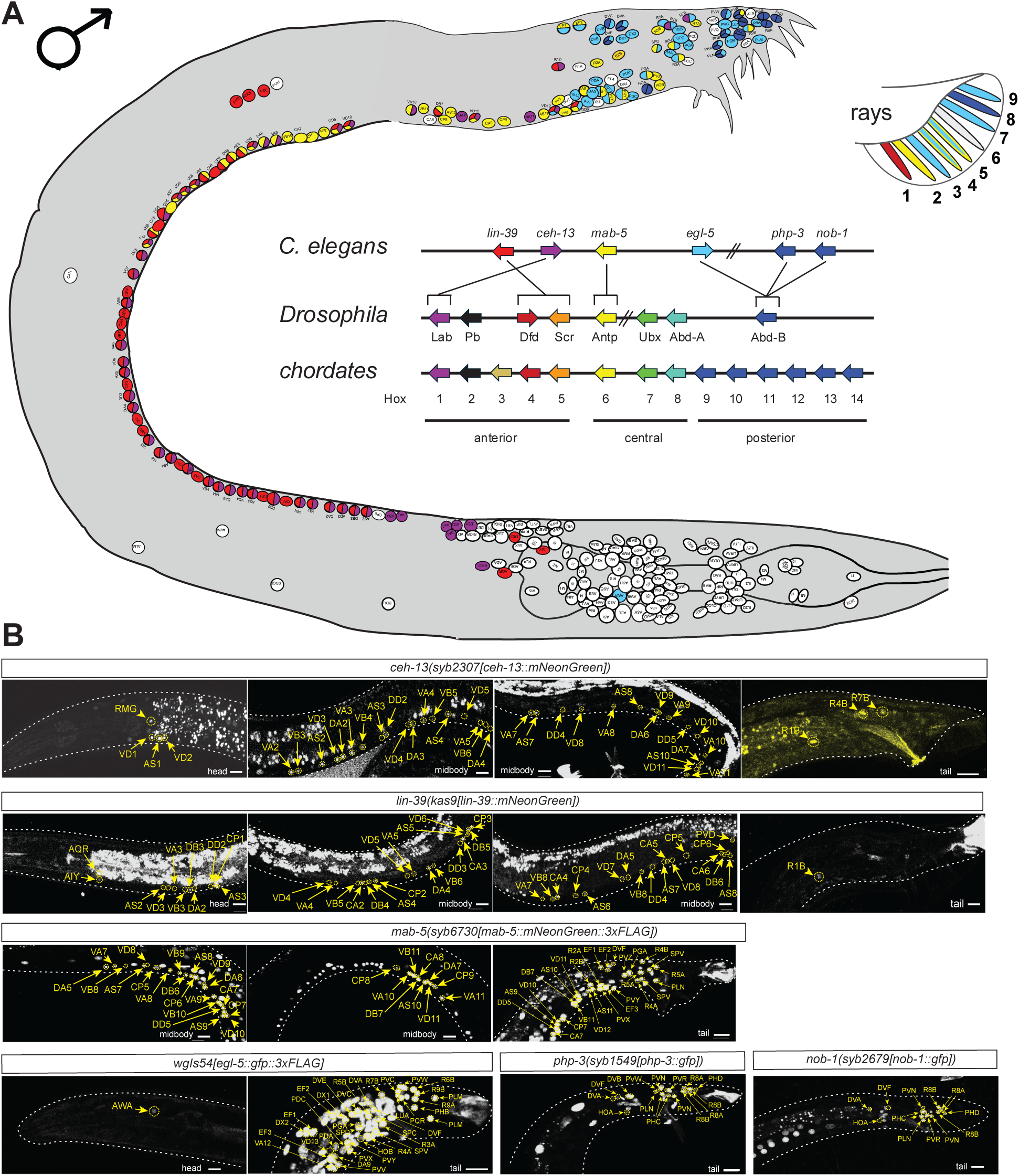
Expression of HOX cluster genes. **A:** Cartoon representation of the expression pattern of HOX genes in male *C. elegans*. Genomic position of HOX genes from *C. elegans*, *Drosophila* and chordates is also represented. HOX cluster gene similarities are from [68]. **B:** Expression of CRISPR/Cas9 engineered reporter alleles and fosmid shows the spatially controlled expression of HOX proteins. Panels show representative images of each protein expression pattern. Circles indicate the HOX protein signal, whilst arrows are used to match the signal to the specific Neural ID. Gut autofluorescence is visible in images for *ceh-13*(*syb2307*) and *lin-39*(*kas9*). Autofluorescence from the male tail is visible in reporters for *lin-39*(*kas9*), *mab-5*(*syb6730*), *egl-5* (reporter array *wgIs54)* and *php-3*(*syb1549*). White dotted lines are used to trace the contours of the animal. To obtain a definitive Neural ID, multiple images acquired with the NeuroPAL landmark (*otIs669* or *otIs696*) are used to obtain a Neuronal ID that is overlaid on representative images. The corresponding images showing overlap with NeuroPAL colors are in **S2 Fig.**

### Temporal dynamics of homeodomain protein expression in the male-specific nervous system

Our expression analysis of all homeodomain proteins has focused on the fully mature nervous system. While we have not examined the onset of expression of homeodomain proteins during developmental specification, we note the presence of widely divergent onsets of expression. Previous work has revealed expression of the AbdB-type EGL-5 HOX already in the epithelial neuroblast precursor of many male-specific neurons at the first larval stage [43], mirrored by neuroblast division defects observed in *egl-5* mutants [47]. On the extreme other end of the spectrum, we find that TTX-1 homeodomain protein only becomes visible in the P12-derived PVX neuron by the third larval stage (**Fig 3A**), even though this neuron is already generated after a few divisions of the P12 neuroblast in the first larval stage [7].

**Fig 3:**
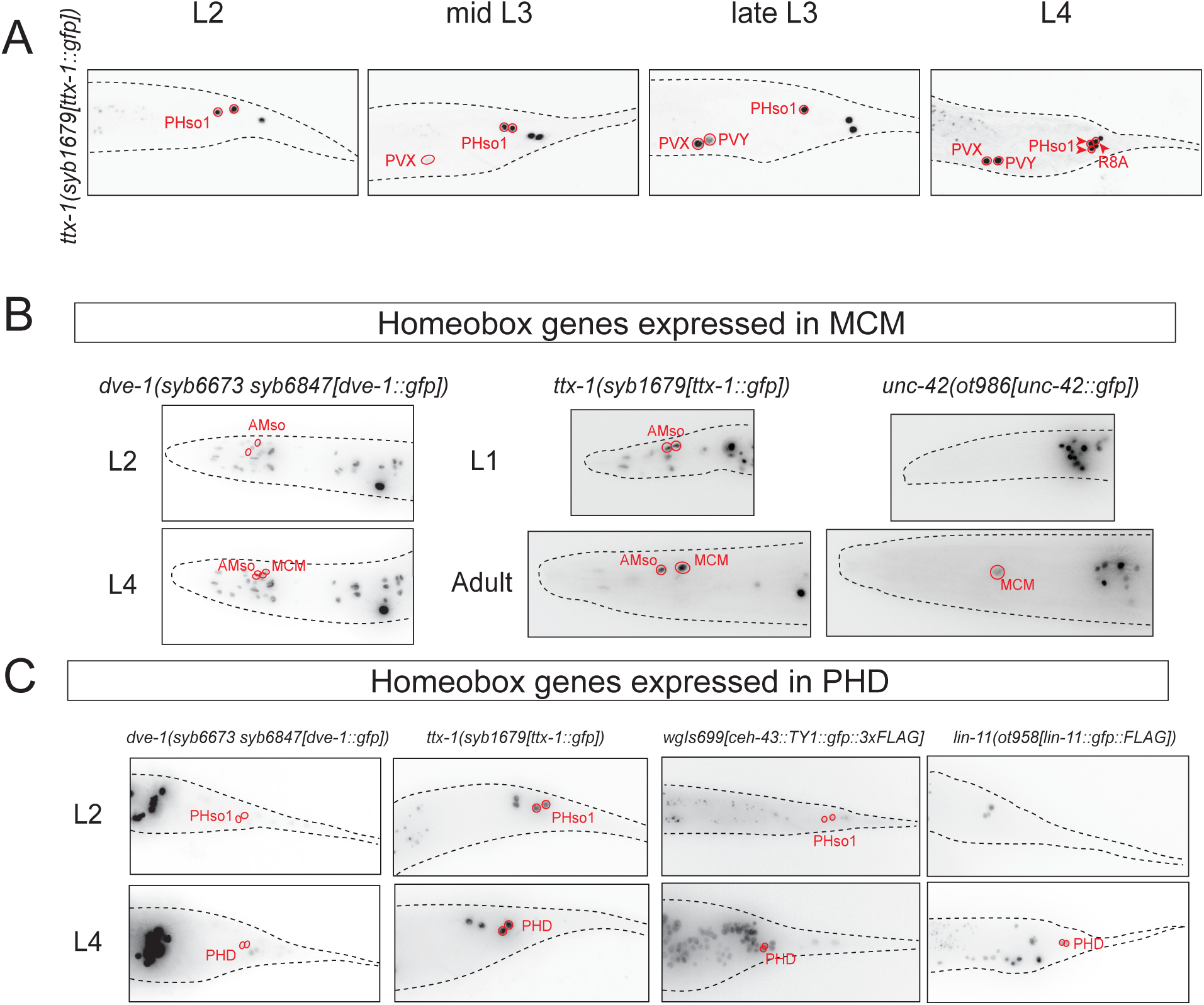
Temporal dynamics of homeobox gene expression in male neurons relative to neuronal transdifferentiation. **A:** Temporal dynamics of *ttx-1(syb1679)* expression in the neurons of the male tail. While PVX is generated in the first larval stage [7], expression of TTX-1 homeodomain protein in PVX is only detectable by the third larval stage. **B:** Temporal dynamics of homeobox genes expressed in MCM. *dve-1(syb6673 syb6847)* and *ttx-1(syb1679)* are expressed in the glial precursors of MCM (AMso) and continue to be expressed in both AMso and MCM after the transdifferentiation event at sexual maturation. *unc-42(ot986)* is expressed only in MCM and not its glial precursor AMso. **C:** Temporal dynamics of homeobox genes expressed in PHD. *dve-1(syb6673 syb6847), ttx-1(syb1679),* and the fosmid-based *ceh-43* reporter *wgIs699* are expressed in the glial precursors of PHD (PHso1) and continue to be expressed in PHD after the transdifferentiation event at sexual maturation. PHso1 to PHD transdifferentiation is direct and does not involve cell division. *lin-11(ot958)* is expressed only in PHD and not in its glial precursor PHso1.

We observed two distinct types of onsets of homeodomain expression in two male-specific neuron classes generated by transdifferentiation. The two male-specific neuron classes, the head interneuron MCM and tail sensory neuron PHD, are generated during sexual maturation through a transdifferentiation process from glial cell types, the AMso glia in the case of MCM and the PHso glia in the case of PHD [10, 11]. We found that a subset of the homeodomain proteins is already expressed in the socket glia from which these neurons are generated: MCM-expressed *dve-1* and *ttx-1* are expressed already in the AMso glia precursor (**Fig 3B**), while PHD-expressed *ceh-43* is expressed already in PHso1 (**Fig 3C**).

Two other homeodomain proteins, *unc-42* and *lin-11*, are, however, only expressed in the respective neuron upon trans-differentiation from glia to neuron: *unc-42* is expressed in MCM, but not AMso (**Fig 3B**) and *lin-11* is expressed in PHD, but not PHso1 (**Fig 3C**).

### A toolbox of terminal differentiation markers for male-specific neurons

One reason why so few previous studies have examined the regulation of differentiation programs in the male-specific nervous system has been the paucity of molecular markers for most male-specific neurons. The construction of the NeuroPAL transgene, which combines a multitude of differentiation markers for the male-specific nervous system [29], as well as our recent mapping of neurotransmitter identities of male-specific neurons [15], has improved the situation, but for many neuron classes still only a limited number of differentiation markers are available [16, 23]. To generate more markers for an analysis of homeobox gene function in the male-specific nervous system, we turned to an analysis of expression of neuropeptide-encoding genes, which display strong and highly neuron type-selective expression throughout the hermaphrodite nervous system, as revealed by previous reporter gene and scRNA analysis [1, 30, 48]. We utilized several previously published neuropeptide reporter alleles, engineered using the CRISPR/Cas9 system and engineered novel, SL2::GFP::H2B-based reporter alleles for a total of 17 neuropeptide-encoding genes, including seven FLP, six NLP and five insulin-like peptides. We analyzed their expression in the male-specific nervous system using the NeuroPAL transgene and discovered highly cell type-specific patterns for each of them, covering in aggregate a large majority of the male-specific nervous system (**Fig 4** and **S3**, **Table 2**). Of particular note is the selective co-expression of four insulin-like genes exclusively in the MCM neuron in the head of adult males (with no expression elsewhere in the male-specific nervous system)(**Fig 4D** and **Table 2**).

**Fig 4:**
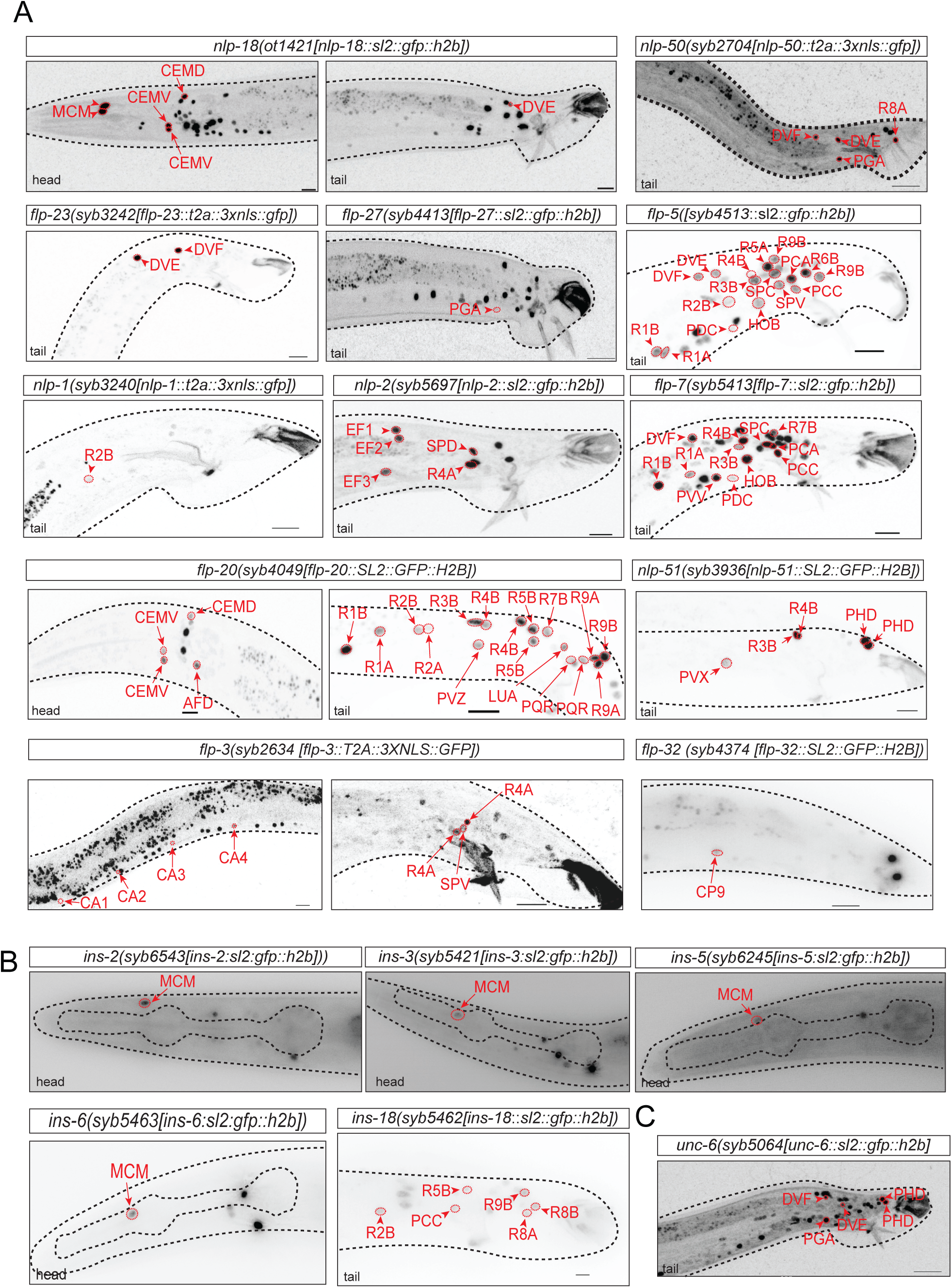
Terminal differentiation markers for male-specific neurons. **A:** CRISPR/Cas9-engineered SL2::GFP::H2B-based reporter alleles reveal previously undescribed expression of several neuropeptides in male-specific neurons. Additional reporter allele data that is not shown here is summarized in **Table 2**. **B:** Expression of CRISPR/Cas9-engineered SL2::GFP::H2B-based reporter alleles of insulin-like peptides in the male nervous system. Four of these, *ins-2(syb6543), ins-3(syb5421), ins-5(syb6245), ins-6(syb5463)*, show expression in MCM but not in other neurons in the male nervous system. *ins-18(syb5462)* is expressed in several neuron types in the male tail. **C:** Stable *unc-6/Netrin* expression in several male tail neurons. CRISPR/Cas9-engineered *unc-6(syb5064)* reporter allele is stably expressed in DVE, DVF, PHD, PGA neurons belonging to distinct ganglia (preanal, lumbar, dorsorectal), suggesting potential function of *unc-6/Netrin* in maintaining neuronal and circuit features. For all panels, the corresponding images showing overlap with NeuroPAL (*otIs669* or *otIs696*) colors are in **S3 Fig.**

In addition to these 17 neuropeptide-encoding genes, we also identified the male-specific sites of expression of *unc-6/Netrin*, whose neuron type-specific expression in select neuron types of the hermaphrodite has served as a valuable cell fate marker for several sex-shared head neuron classes [49](**Fig 4C**, **Table 2**). We found that UNC-6/Netrin is selectively expressed in four male-specific neuron classes (DVE, DVF, PHD, PGA), covering three different tail ganglia (preanal, lumbar, dorsorectal). Expression is observed not just transiently, as one would perhaps expect from a gene so prominently involved in axon pathfinding [50], but is rather stably observed after males have reached maturity. Such maintained expression is suggestive of UNC-6/Netrin function in maintaining neuronal features, such as proper synaptic connectivity.

### The Pitx-homolog *unc-30* is required for PGA interneuron differentiation

Armed with these molecular markers, we set out to examine whether homeobox genes affect neuronal differentiation to an extent similar to what has been observed in other neuronal contexts in the hermaphrodite. We first examined one of the two most sparsely expressed homeobox genes, the *C. elegans* homolog of the vertebrate Pitx genes, *unc-30,* which, in the context of the male-specific nervous system, is exclusively expressed in a single male-specific neuron class, the PGA interneuron in the preanal ganglion. PGA is one of a total of 4 pre-anal ganglion interneurons generated by the P11 epidermal blast cells in response to Wnt and EGF signaling [51]. The terminally differentiated state of PGA is one of the few neurons in the *C. elegans* nervous system that expresses multiple neurotransmitter systems: Based on *unc-17/VAChT* expression, PGA is cholinergic but also uses an additional, unknown neurotransmitter, based on the expression of the GABA/Glycine vesicular transporter *unc-47/VGAT* and the concomitant absence of *unc-25/GAD* [15]. Moreover, PGA re-uptakes and re-utilizes serotonin, as inferred from absence of *tph-1/TPH* expression, 5HT antibody staining, *cat-1/VMAT* expression and *mod-5/SERT* expression [15]. In addition, our marker analysis described above identified two neuropeptides (*flp-27* and *nlp-50)* expressed in PGA, as well as the axon guidance/synaptogenic UNC-6/Netrin protein (**Fig 4A,C**). PGA is also marked by a specific color-code in NeuroPAL [29].

To assess *unc-30* function, we engineered a molecular null allele *(ot1186)* using CRISPR/Cas9 genome engineering [35]. We find that the entire cohort of neurotransmitter and neuropeptide reporters fails to be properly expressed in the PGA neuron of these *unc-30* null mutants (**Fig 5A** and **5B**). Moreover, 5HT antibody staining, which we find to be *mod-5/SERT-*dependent, as expected from absence of 5HT biosynthesis pathway genes, is absent in *unc-30* null mutants (**Fig 5C**). *unc-6/Netrin* reporter allele expression is also affected and so is the normal color code of NeuroPAL (**Fig 5A**). However, since the panneuronal marker of the NeuroPAL transgene is not affected in PGA, we can conclude that PGA is generated in *unc-30(ot1186)* mutant animals but fails to adopt its proper identity. *unc-30* therefore classifies as a terminal selector of PGA identity, mirroring its terminal selector function in several sex-shared neuron classes previously examined in the hermaphrodite, namely, the D-type motorneuron, the PVP and the AVJ neuron classes [5, 22, 52, 53]. Intriguingly, the NeuroPAL transgene generates a novel TagBFP signal in PGA in *unc-30(ot1186)*(**Fig 5A**), indicating that PGA may have undergone an identity transformation to another neuron class. We cannot presently tell what this alternative neuronal identity may be since the NeuroPAL transgene expresses TagBFP from 12 different promoters, expressed in 10 different neuron classes [29].

**Fig 5:**
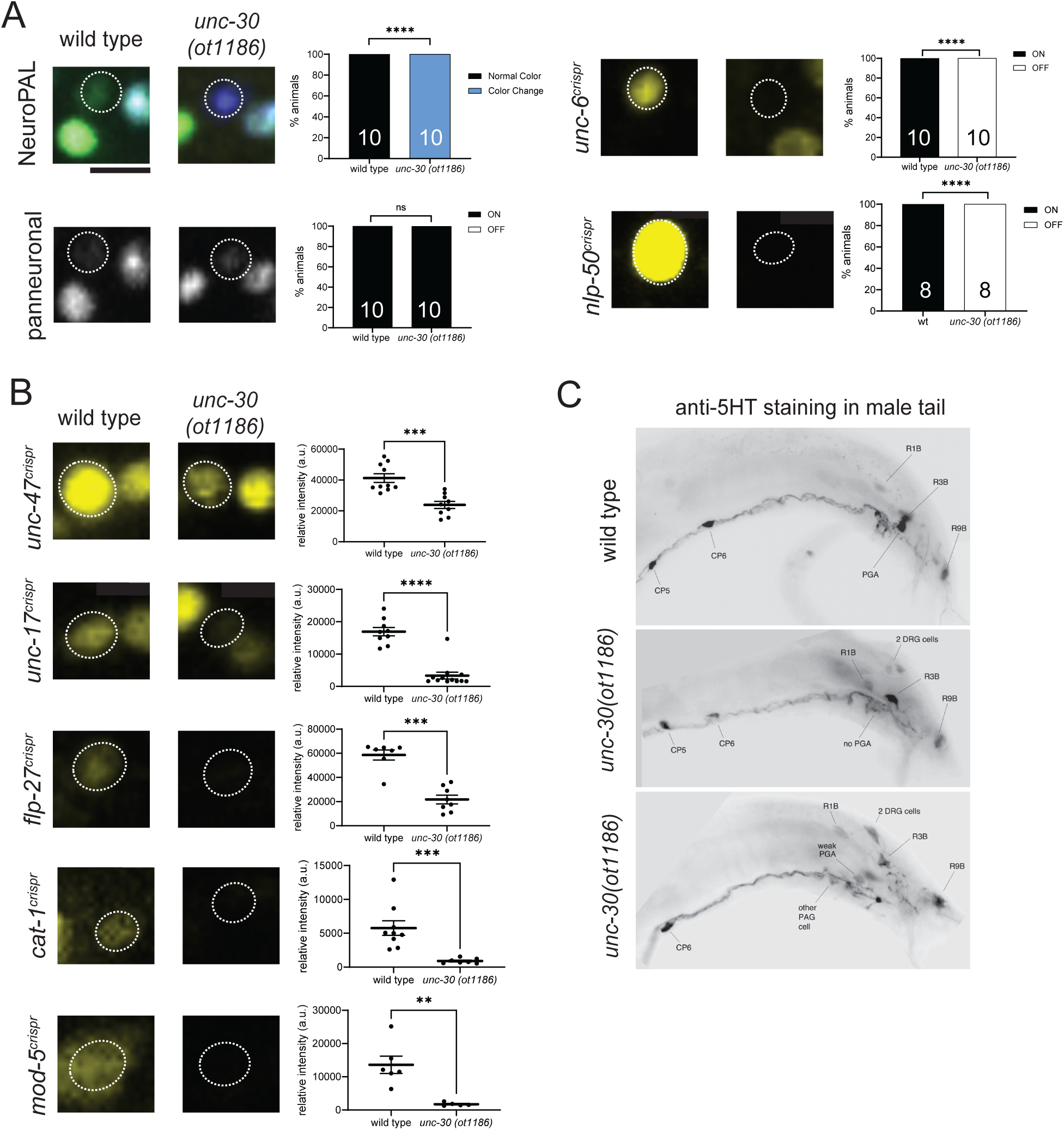
The *unc-30* Pitx-type homeobox gene affects differentiation of the PGA neuron. **A**: In *unc-30(ot1186)* null mutant animals, the NeuroPAL (*otIs669*) color code of PGA is changed from orange to blue. Expression of *unc-6(syb5064)* and *nlp-50(syb2704)* in PGA is abolished in *unc-30(ot1186)* null mutants. Number of animals scored are shown within each bar. **B:** *unc-30(ot1186)* null mutants show decreased expression of several PGA markers, including *unc-47(syb7566), unc-17(syb4491), flp-27(syb4413), cat-1(syb6486),* and *mod-5(vlc47)*. Statistical analyses for panel A were performed using Fischer’s exact test. For panel B, the Mann-Whitney test was used. Error bars indicate SEM. ****p ≤ 0.0001, ***p ≤ 0.001, **p ≤ 0.01. **C:** Loss of anti-serotonin antibody staining in *unc-30(ot1186)* null mutant animals. 0/52 male tails had strong PGA staining; 3/15 had a weakly staining PGA (bottom panel).

### The Prop1-homolog *unc-42* and the Otx-homolog *ttx-1* control different aspects of MCM neuron differentiation

We discovered a similar terminal selector role for the Prop1 homolog *unc-42* homeobox gene, which, like *unc-30*, is expressed in a single male-specific neuron class, the peptidergic MCM neuron in the head of the worm, after it has transdifferentiated from the AMso cell (**Fig 3B**). Our analysis of neuropeptide reporter alleles defined four markers for MCM, including reporter alleles for *pdf-1, ins-2, ins-5* and *ins-6.* We found that all four markers fail to be properly expressed in the MCM neurons of *unc-42* null mutant animals, containing an entire deletion of the *unc-42* coding region, generated by CRISPR/Cas9 genome engineering (**Fig 6A and Methods**). Expression of the panneuronal *rab-3* marker is unaffected in *unc-42* null mutant animals, suggesting that MCM neurons are properly generated (i.e. transdifferentiated from the AMso amphid socket glia), but fail to adopt their unique identity (**Fig 6A**). We conclude that *unc-42/Prop1* acts as a terminal selector of MCM identity.

**Fig 6:**
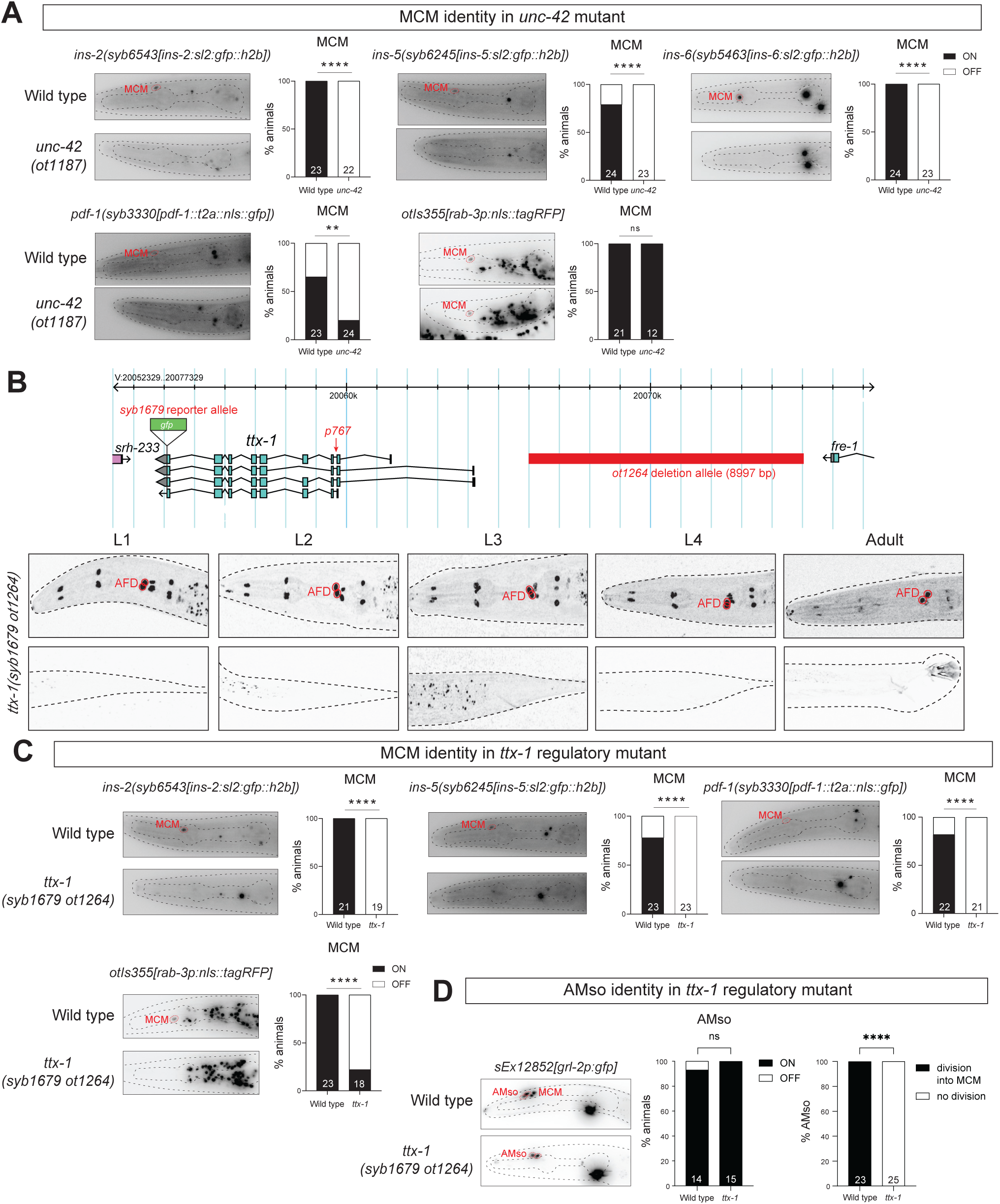
Effect of the Otx-type *ttx-1* and Prop-type *unc-42* homeobox gene on MCM neuron differentiation. **A:** MCM terminal identity markers, but not a pan-neural marker, are lost in *unc-42(ot1187)* null mutants. Gray-scale images are representative of expression of 4 terminal identity markers [*ins-2(syb6543), ins-5(syb6245), ins-6(syb5463)* and *pdf-1(syb3330)*] and one pan-neuronal marker (*rab-3* array *otIs355)* in a wild type (control) and *unc-42(ot1187)* null mutant background. Dotted lines are used to trace the contours of the animal and its pharynx. Bar graphs show the percentage of animals that retained or lost expression in MCM. **B:** A *gfp-*tagged, *cis-*regulatory allele of *ttx-1, syb1679 ot1264,* results in loss of expression in male-specific neurons MCM and PHD. Cartoon representation of the *ttx-1* locus showing the extent of the *ot1264* deletion (red bar) and the *syb1679* edit which inserts a gfp in frame with *ttx-1* for visualizing the protein expression. Representative images show that in a *ot1264* deletion background, *ttx-1* is not expressed in MCM and PHD. Dotted lines are used to trace the contours of the animal. **C:** MCM terminal identity markers and a pan-neural marker, are lost in *ttx-1(syb1679 ot1264) cis-*regulatory mutants. Gray-scale images are representative of expression of 3 terminal identity markers (*ins-2(syb6543), ins-5(syb6245)* and *pdf-1(syb3330)* reporter alleles) and one pan-neuronal marker (*rab-3* reporter array *otIs355*) in a wild type (control) and *ttx-1(syb1679 ot1264)* mutant background. Dotted lines are used to trace the contours of the animal and its pharynx. Bright extra signal between the first and second bulb of the pharynx in *ttx-1*(*syb1679 ot1264*) mutant animals is likely TTX-1::GFP (See Fig 6B, syb1679) signal coming from AFD. The *ot1264* regulatory allele does not disrupt *ttx-1* expression in a number of cells, including AFD [5]. Bar graphs show the percentage of animals that retained or lost expression in MCM. **D:** *ttx-1(syb1679 ot1264)* animals show impaired MCM differentiation from AMso glia. Gray-scale images are representative of expression of *grl-2* (reporter array *sEx12853*), an AMso terminal identity marker in a wild type (Control) and *ttx-1(syb1679 ot1264)* mutant background. Dotted lines are used to trace the contours of the animal and its pharynx. Left bar graph shows the percentage of animals that retained or lost *grl-2 (*reporter array *sEx12853)* expression in AMso. Right bar graph shows the percentage of AMso that gave rise to MCM. During AMso to MCM transdifferentiation in wild type animals, the GFP protein from *grl-2* expressing AMso perdures into MCM as it divides [10]. In *ttx-1(syb1679 ot1264)* mutants, no perdurance is seen. Statistics: **p value ≤0.01, ****p value ≤0.0001, ns, not significant. Testing was performed using the Fisher’s Exact test.

Since homeodomain proteins are known to act in combinations, we also tested the role of another homeobox gene that we found to be expressed in MCM from their initial differentiation throughout adulthood, the Otx-type homeobox gene *ttx-1* (**Fig 3B**). To analyze *ttx-1* function, we could not utilize a molecular null allele of the entire locus since such a deletion results in embryonic lethality [5]. In a previous study on *ttx-1* function in the hermaphrodite nervous system, we had circumvented this problem by generating a *cis-* regulatory allele, *ot1264*, a 9 kb deletion of an enhancer region that resulted in loss of *ttx-1* expression in several sex-shared neurons, but not in other tissues in which *ttx-1* function is required for embryonic viability [5]. Using a *gfp-*tagged *ttx-1* reporter allele, we examined whether this *cis*-regulatory allele also eliminates expression of *ttx-1* from the male-specific neuron classes in which *ttx-1* is normally expressed in. We found this to indeed be the case (**Fig 6B**). Using this *cis*-regulatory allele, we found that elimination of *ttx-1* displays the same phenotype as *unc-42* null mutant animals: expression of *pdf-1, ins-2* and *ins-5* is lost (**Fig 6C**). In this case, however, we also failed to detect expression of the *rab-3* panneuronal marker, indicating that the MCM neurons may not be properly generated at all (**Fig 6C**).

Since *ttx-1* is, in contrast to *unc-42*, already expressed in the AMso glia cell from which the MCM neuron transdifferentiated (**Fig 3B**), we assessed AMso glia (as well as PHso) differentiation defects in *ttx-1(syb1679 ot1264)* mutants. We found no effect on the expression of the *grl-2* glia marker in juvenile AMso glia (**Fig 6D**). However, we noted that after the cell division of the embryonically generated AMso during sexual maturation in larval stage animals, which generates the adult AMso and the MCM neurons, the expression of *grl-2* from embryonically generated AMso perdures in the MCM neurons [10]. We fail to detect such perdurance in *ttx-1(syb1679 ot1264)* mutants (**Fig 6D**). Taken together with the lack of *rab-3* expression where we expect MCM to be, we conclude that in the absence of *ttx-1*, the MCM neuron fails to be generated, possibly due to a failure of embryonically generated AMso to divide upon sexual maturation.

### The LIM homeobox gene *lim-6* affects DVE and DVF differentiation

The LIM homeobox gene *lim-6,* the *C. elegans* ortholog of vertebrate Lmx1/2, is known to control the differentiation program of the sex-shared DVB neuron, located in the dorsorectal ganglion [54, 55]. The expression of *lim-6* in two other, male-specific neurons of the dorsorectal ganglion, DVE and DVF, as well as the existence of cell markers for these two neurons, prompted us to assess the effect of *lim-6* on the differentiation of these neurons. We found that the characteristic color code of the NeuroPAL transgene in the DVE and DVF neurons of *lim-6* null mutants is defective (**Fig 7A**). DVE and DVF do not synthesize a currently known fast neurotransmitter system, but express the vesicular neurotransmitter transporter *unc-47/VGAT* [15] and this expression is significantly affected upon loss of *lim-6* (**Fig 7B**). Our neuropeptide expression analysis identified three neuropeptides expressed in these neurons, *flp-23* and *nlp-50* (expressed in both DVE and DVF) and *nlp-18* (expressed only in DVE)(**Fig 4**). We found that expression of *nlp-18* in DVE is affected in *lim-6* null mutants (**Fig 7B**), while there is a reduction in the expression levels of *flp-23* in DVE but not DVF. However, expression of *nlp-50* in both neurons is not affected in *lim-6* null mutants (**Fig 7C**). We conclude that *lim-6* is required for the proper differentiation of DVE and DVF.

**Fig 7:**
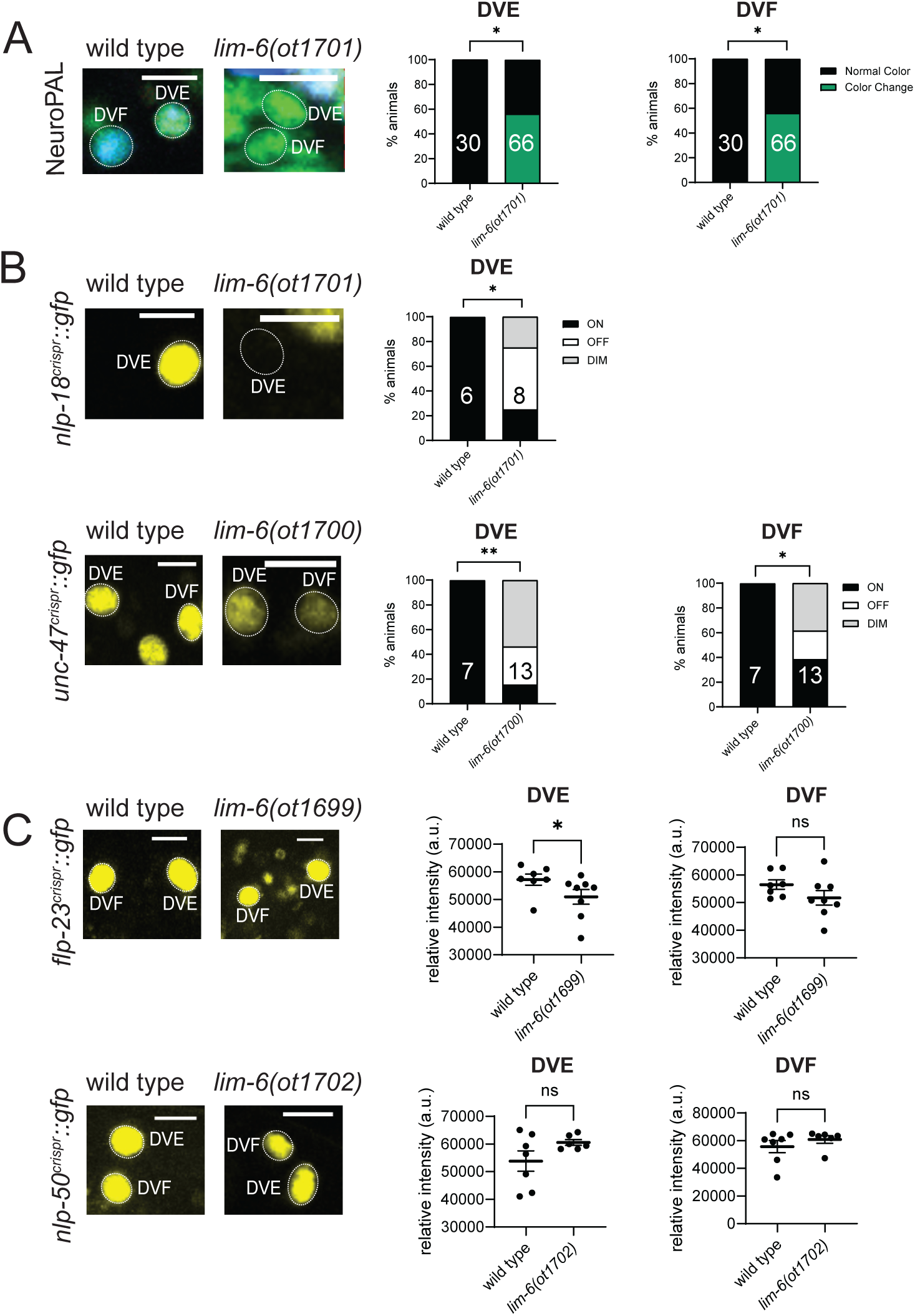
The Lmx-type *lim-6* homeobox gene affects differentiation of DVE and DVF neurons. **A:** In nearly half of *lim-6(ot1701)* null mutant animals, the NeuroPAL (*otIs696*) colors of DVE and DVF are changed from turquoise to green, suggesting loss of blue-expressing reporters. *lim-6* null mutants exhibit low mating efficiency. In lieu of crosses, molecularly identical null deletions were generated in the background of the markers in panels B and C by CRISPR/Cas9-engineering using the same guide RNAs and repair templates (see Methods). These alleles are *lim-6(ot1699)*, *lim-6(ot1700), lim-6(ot1701)* and *lim-6(ot1702)*. **B:** *lim-6(ot1701)* and *lim-6(ot1700)* null mutants display defects in the expression of DVE and DVF markers *nlp-18(ot1421)* and *unc-47(syb7566)*. **C:** *lim-6(ot1699)* and *lim-6(ot1702)* null mutants do not show defects in the expression of the DVE and DVF markers *nlp-50(syb2704)* and *flp-23(syb3242)*. Statistical analyses for panels A and B were performed using Fischer’s exact test. For panel C, the Mann-Whitney test was used. Error bars indicate SEM. **p ≤ 0.01, *p ≤ 0.05, ns not significant.

The DVE and DVF neurons also co-express the Eve-homolog *vab-7*. We found that in *vab-7* mutants, marker gene expression in neurons of the dorsorectal ganglion do not appear to be affected, but we noted an increased number of neurons in the ganglion, indicating lineage division defects, which we did not pursue further.

### The LIM homeobox gene *lin-11* has a range of effects on several neuron classes

We extended our homeobox mutant analysis to additional neuron types and found that removal of the LIM homeobox gene *lin-11,* the *C. elegans* ortholog of vertebrate LHX1/5, has complex, differential effects on the differentiation of several different male-specific neuron classes. The PHD neurons, generated during sexual maturation by transdifferentiation from PHso1 glia, are normally generated in *lin-11* null mutants and still express both their cholinergic identity (*unc-17/VAChT),*as well as the single Ig domain protein *oig-8* (**Fig 8A**), a previously described marker of PHD identity [11, 56]. However, the NeuroPAL color code, as well as expression of the *nlp-51* neuropeptide and *unc-6/*Netrin are defective (**Fig 8A and S4A**). The Otx-type homeobox gene *ttx-1,* which is also expressed in PHD, also affects *unc-6/Netrin* expression and the NeuroPAL color code, but does not affect *nlp-51*, *oig-8* or PHD’s cholinergic identity (*unc-17/VAChT*) (**Fig 8B** and **S4B**). *lin-11; ttx-1* double mutants do not show more severe defects in PHD differentiation than each single mutant, i.e. neither *oig-8* nor *unc-17/VAChT* markers are affected (**Fig 8C**).

**Fig 8:**
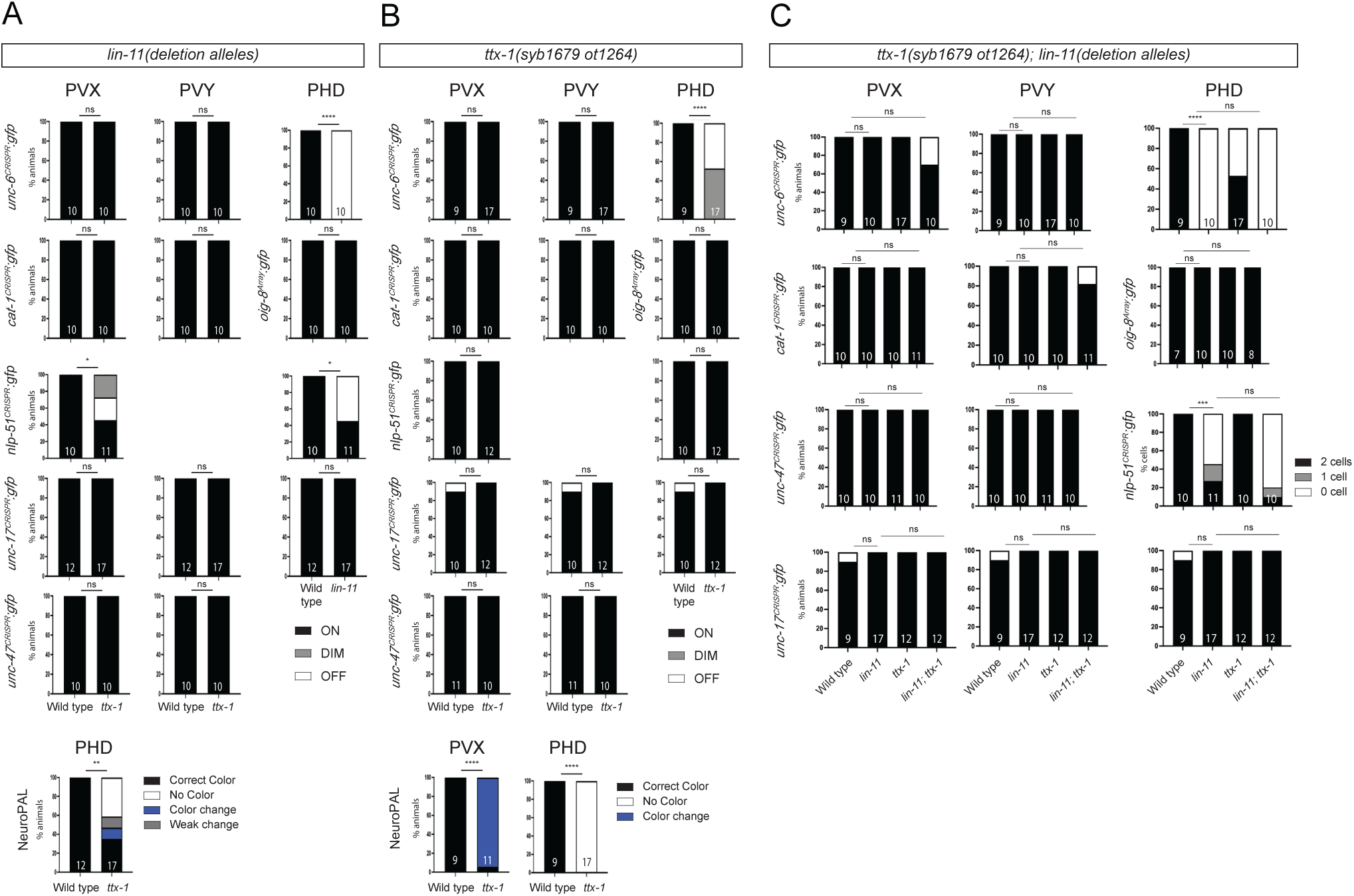
Effect of the LIM-type homeobox gene *lin-11* and the Otx-type *ttx-1* homeobox gene on PVX, PVY, PHD differentiation. **A:** Bar graphs show the percentage of animals that retained or lost signal for 6 terminal identity markers (*unc-6(syb5064), cat-1(syb6486), oig-8* array*(drpIs4), nlp-51(syb3936), unc-17(syb4491), unc-47(syb7566)*) as well as quantifying the NeuroPAL(*otIs669*) color in the male-specific neurons (PVX, PVY and PHD) of wild type and *ttx-1(syb1679 ot1264)* regulatory mutants **B:** Bar graphs show the percentage of animals that retained or lost signal for 6 terminal identity markers (*unc-6(syb5064), cat-1(syb6486), oig-8* array*(drpIs4), nlp-51(syb3936), unc-17(syb4491), unc-47(syb7566)*) as well as quantifying the NeuroPAL(*otIs669*) color in the male-specific neuron (PVX, PVY and PHD) of wild type and *lin-11* null mutants *(lin-11(ot1483);unc-6(syb5064), lin-11(ot1472);cat-1(syb6486), lin-11(ot1521);oig-8* array *(drpIs4), lin-11(ot1444);nlp-51(syb3936), lin-11(ot1026);unc-17(syb4491), lin-11(ot1497);unc-47(syb7566))*. Using CRISPR/Cas9 and HDR multiple but genetically identical null alleles of *lin-11* were obtained (See Methods). **C:** Bar graphs show the percentage of animals that retained or lost signal for 6 terminal identity markers (*unc-6(syb5064), cat-1(syb6486), oig-8* array *(drpIs4), nlp-51(syb3936), unc-17(syb4491), unc-47(syb7566)*) in the male-specific neurons (PVX, PVY and PHD) of wild type and *ttx-1(syb1679 ot1264);lin-11* double mutants *(ttx-1(syb1679 ot1264);lin-11(ot1483);unc-6(syb5064), ttx-1(syb1679 ot1264);lin-11(ot1472);cat-1(syb6486), ttx-1(syb1679 ot1264);lin-11(ot1521);oig-8* array *(drpIs4), ttx-1(syb1679 ot1264);lin-11(ot1444);nlp-51(syb3936), ttx-1(syb1679 ot1264);lin-11(ot1026);unc-17(syb4491), ttx-1(syb1679 ot1264);lin-11(ot1497);unc-47(syb7566))*. Using CRISPR/Cas9 and HDR multiple but genetically identical null alleles of *lin-11* were obtained (See Methods). Statistics: *p value ≤0.05, **p value ≤0.01, ***p value ≤0.001, ****p value ≤0.0001, ns, not significant. Testing was performed using the Fisher Exact test.

Besides PHD, the *lin-11* and *ttx-1* homeobox genes are also co-expressed in the PVX and PVY neurons. We did not observe any differentiation defects in PVX or PVY in *lin-11, ttx-1* or *lin-11; ttx-1* double mutants based on intact *unc-6/Netrin* and *oig-8* expression, aminergic identity (*cat-1/VMAT*), cholinergic identity (*unc-17/VAChT*) and other terminal identity features (*unc-47/VGAT*) (**Fig 8A-C**). However, we found that in *lin-11* null mutant animals, but not in *ttx-1* mutants, the expression of *nlp-51* was affected in PVX neurons (**Fig 8A and S4A**). *ttx-1* may nevertheless have a partial impact on PVX differentiation since we observed a change in the NeuroPAL color code (**Fig 8B** and **S4B**).

In the ventral nerve cord, *lin-11* is expressed in a subset of CA and all CP neurons. Each of these neuron classes falls into distinct subtypes based on neurotransmitter expression [15], neuropeptide expression (**Table 2**) and color codes of NeuroPAL transgene [29]. We found that cholinergic identity (*unc-17/VAChT* expression) of the CA1-4 neurons, which normally express *lin-11,* is unaffected in *lin-11* null mutants (13/14 animals show normal expression of an *unc-17/VAChT* reporter allele). Instead, several of the CP neurons (CP0, CP5, CP6, CP7, CP8), as well as CA7, which are either not cholinergic or express *unc-17* only very weakly, express *unc-17/VAChT* either ectopically or much more strongly in *lin-11* null mutants (**Fig 9A** and B). Ectopic expression in *lin-11* mutants was not limited to *unc-17/VAChT* since we observed ectopic expression of *cat-1/VMAT* in CP8 as well (**Fig 9C**). Moreover, there is a concomitant novel NeuroPAL color code (gain of blue color) observed in the CP1-6 neurons of *lin-11* mutants (**Fig 9D**). Since the CA neurons, which are sisters of the CP neurons, are cholinergic [15], a CP to CA identity switch could be envisioned. However, such transformation is unlikely since first, the change in NeuroPAL color code is not consistent with such a switch (NeuroPAL does not mark the CA neurons with any signal beyond the panneuronal signal, yet there is a gain in orange signal in *lin-11* mutants) and, second, we find 5HT antibody staining of CP neurons to be unaffected in *lin-11* mutants (52/55 animals show normal 5HT staining), indicating the CPs do retain aspects of their original identity. At this point, we can only infer that in *lin-11* mutants, the CP neurons are not properly specified.

**Fig 9:**
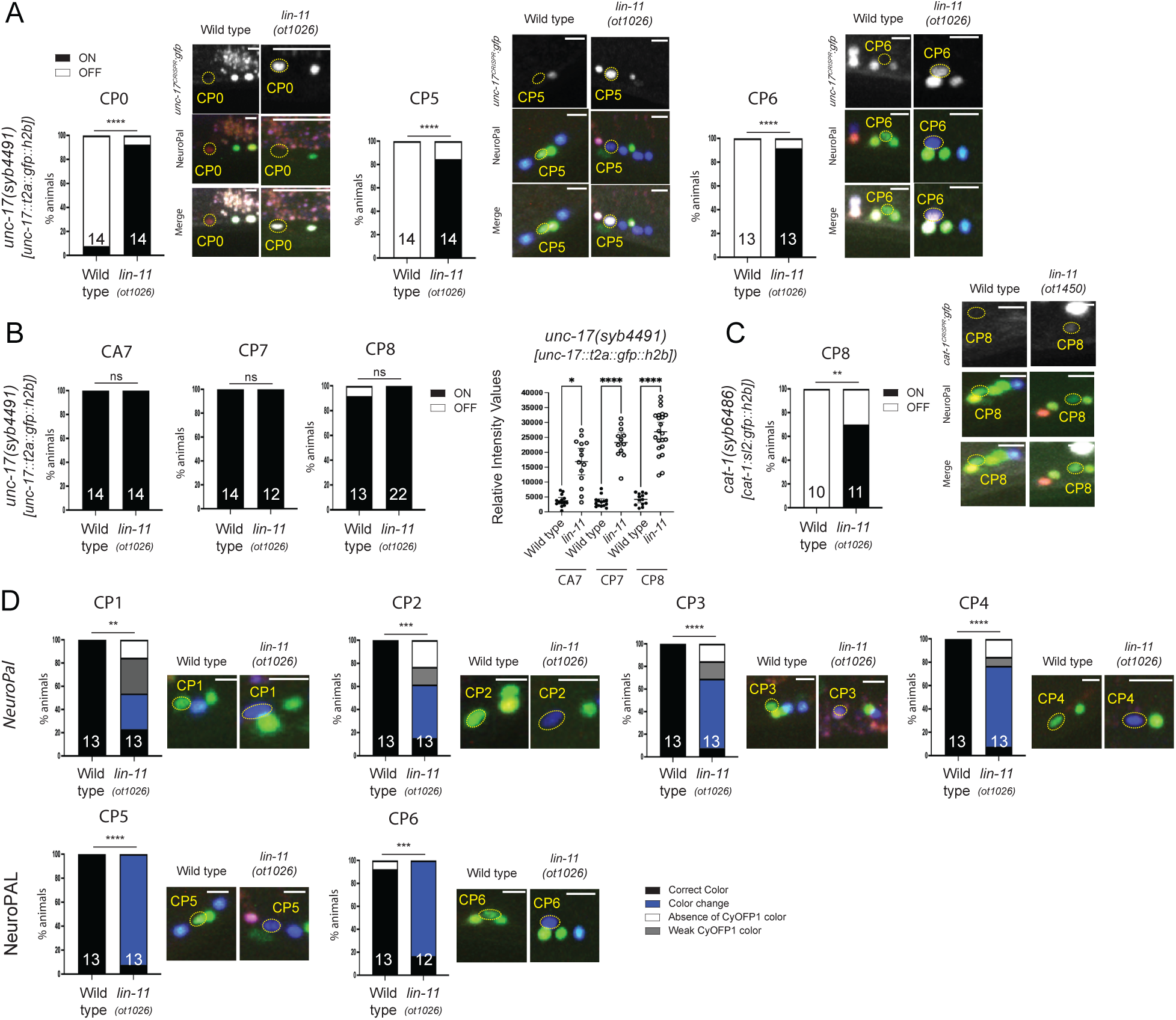
Effect of the LIM homeobox gene *lin-11* on CP neuron differentiation. **A:** Extra *unc-17* positive CP neurons are found in *lin-11(ot1026)* null mutants. Representative images show the expression of *unc-17(syb4491)*, NeuroPAL*(otIs669)* landmark and their merge in wild type and *lin-11(ot1026)* mutants. Bar graphs show the percentage of animals expressing ectopic *unc-17*. **B:** Absence of *lin-11* increases the expression of *unc-17* in several CP neurons. Bar graphs show the percentage of animals expressing *unc-17* as well as intensity values in wild type and *lin-11(ot1026)* null mutants. **C:** *lin-11(ot1026)* mutants show alter NeuroPAL colors in CP neurons. Representative images show an overlay of the NeuroPAL*(otIs669)* pseudocolors in wild type and *lin-11(ot1026)* null mutants. CP neurons have a stronger blue fluorophore signal in *lin-11(ot1026)* null mutants compared to wild type. Bar graphs show the percentage of animals with no color change (Correct Color), a color change (Color Change), absence of the orange color (Absence of CyOFP1 color) and a dim orange color (Weak CyOFP1 color). **D:** *lin-11* null mutants show ectopic *cat-1* expression in CP8 male-specific neurons. Representative images show the expression of *cat-1(syb6486)*, NeuroPAL*(otIs669)* landmark and their merge in wild type and *lin-11(ot1450)* mutants. Bar graphs show the percentage of animals expressing ectopic *cat-1*. Statistics: *p value ≤0.05, **p value ≤0.01, ***p value ≤0.001, ****p value ≤0.0001, ns, not significant. Testing was performed using the Fisher Exact test and Student t-test for quantification of Relative Intensity Values

### Behavioral deficits of *lin-11* and *ttx-1* mutants

Due to their strong locomotory defects, we were not able to assess the behavioral consequences of loss of MCM differentiation in *unc-42/Prop1* null mutants or loss of PGA differentiation in *unc-30/Pitx* null mutants. However, superficially normal locomotion of *lin-11(n389)* animals and *ttx-1(syb1679 ot1264)* animals did allow us to assess possible effects on PHD, PVX and/or PVY function. These three neurons are among the very few male-specific neurons to which functions were previously assigned. Specifically, the PVX and PVY interneurons have been shown to be involved in the male’s contact-based scanning of the hermaphrodite’s surface for the vulva [34]. This scanning behavior also requires the PHD neuron, which is presynaptic to both PVX and PVY [11]. The expression of *lin-11* and *ttx-1* in PHD, PVX and PVY, combined with the observed effect of these transcription factors on a subset of differentiation markers in at least two of these neurons, prompted us to examine male mating behavior in *lin-11* and *ttx-1* mutant males. We found that both tail contact and scanning behavior of *ttx-1* mutant males is indeed defective, as predicted by *ttx-1’s* expression in PVX, PVY and PHD (**Fig 10A**). Scanning behavior, but not tail contact, is also defective in *lin-11* mutant males (**Fig 10B**). PHD is also involved in another aspect of the mating behavior, the Molina maneuver, a re-engagement process of males with a mate after an unsuccessful mating attempt [11]. Molina maneuvers are defective in *lin-11* mutant males, but not in *ttx-1* mutants, consistent with *lin-11* mutant males appearing to have a more severe PHD differentiation defect than *ttx-1* mutant males (**Fig 8A** and **B**).

**Fig 10:**
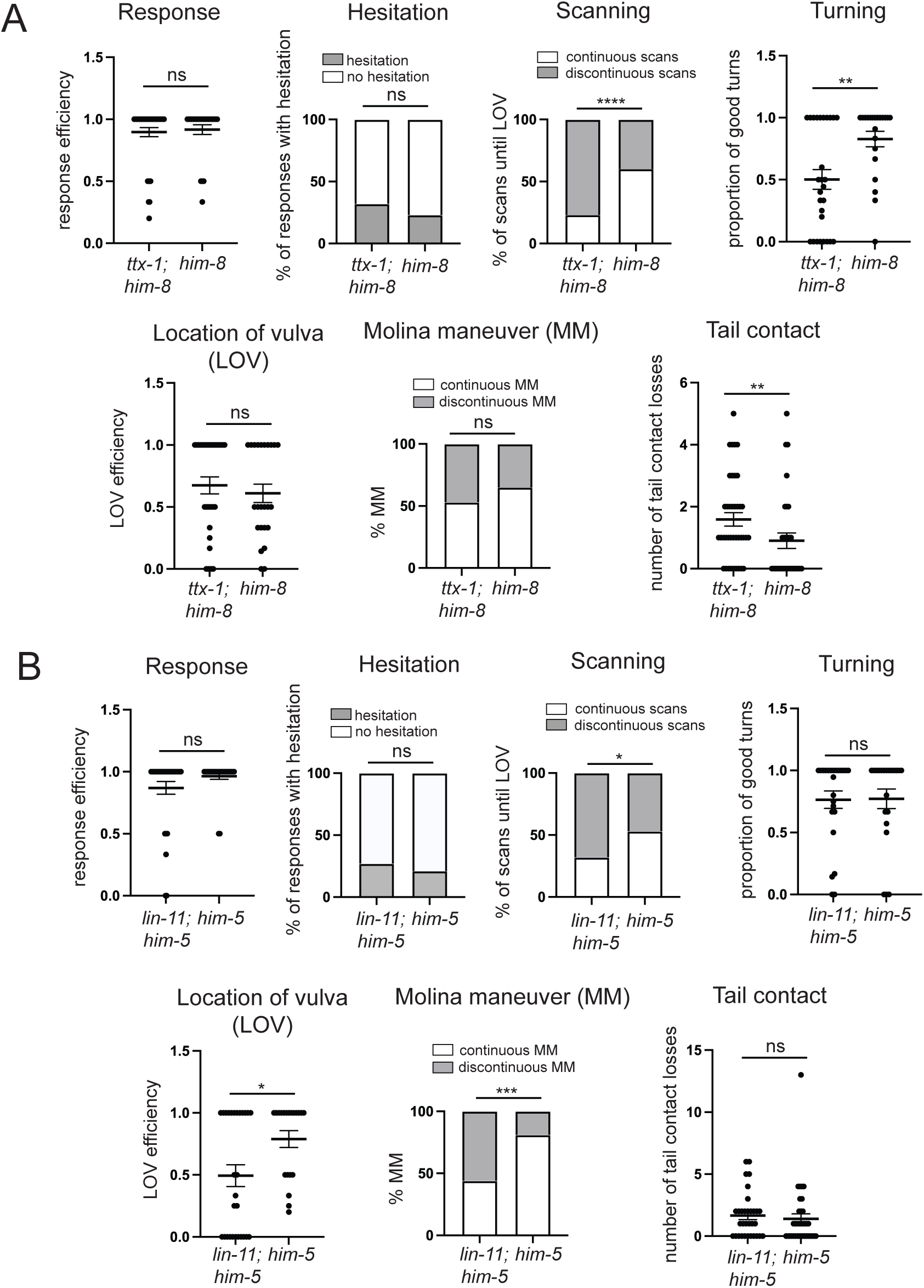
*lin-11* and *ttx-1* mutants display male mating defects. **A:** Male mating steps in *ttx-1(syb1679 ot1264); him-8(e1489) IV* mutant males, mated with *unc-51* mutant hermaphrodites. See Methods for details on scored behavior. Mann-Whitney tests were used to compare efficiencies of response, turning, location of vulva and tail contact of control and mutant males; χ^2^ was used to compare the proportion of responses with hesitation and proportion of continuous scans and Molina Maneuvers. n.s = not significantly different, p≥0.05, ** p≤ 0.01, **** p≤0.0001, n= number of events; 23 *ttx-1* and 28 *him-8* males were scored. **B:** Male mating steps in *lin-11(n389) I; him-5(e1467)* mutant males, mated with *unc-51* mutant hermaphrodites. See Methods for details on scored behavior. Mann-Whitney test was used to compare efficiencies of response, turning, location of vulva and tail contact of control and mutant males; χ^2^ was used to compare the proportion of responses with hesitation and proportion of continuous scans and Molina Maneuvers. n.s = not significantly different, p≥0.05, * p≤ 0.05, *** p≤0.001, n= number of events; 18 *lin-11* and 20 *him-5* were scored.

While other aspects of male mating behavior are intact in *ttx-1* and *lin-11* mutants, we also noted two defects that are not predicted by their expression pattern (**Fig 10A** and **B**): *lin-11* mutants display defects in vulval stop behavior and *ttx-1* mutant display defects in turning behavior. Whether these defects are also the result of PVX, PVY or PHD differentiation defects is currently unknown.

## DISCUSSION

### The male-specific nervous system of *C. elegans*

The nervous system of male nematodes is substantially larger than the nervous system of hermaphrodites, owing largely to sex-specific neuroblast proliferation that generate a wide spectrum of distinct neuron types only in males [7]. While brain size differences are also apparent in different areas of human male and female brains [57], the underlying cellular basis of such differences remains unclear in mammals. While the number, lineage and morphology of male-specific neurons in *C. elegans* have been precisely delineated [7, 9], mechanisms that specify the unique identities of these male-specific neurons have mostly been explored in the context of developmental patterning, namely of (a) the ray sensory neuron classes, a prominent subgroup of male-specific neurons [16, 25, 58, 59], (b) two male-specific neuron classes that are generated via transdifferentiation (MCM, PHD)[10, 11], (c) one male-specific neuron class whose sex-specificity is the result of sex-specific cell death (CEM) [20, 21, 60] and (d) several male-specific motor and interneurons in the ventral cord (CA,CP)[23, 24]. Earlier patterning information is available for the developmental specification programs of other male-specific neurons [61], yet information about terminal differentiation programs had been limited. We have begun here to rectify these limitations, by (a) establishing a protein expression atlas of a family of transcription factors, homeodomain proteins, and (b) defining the role of these proteins in cell fate specification using a new set of reagents that allowed us to define cell identity.

### Homeobox gene expression

In the hermaphrodite, an analysis of homeodomain protein expression has shown that each individual terminally differentiated neuron class can be defined by a unique combination of homeodomain proteins [4]. Even though we analyzed only half of all conserved *C. elegans* homeodomain proteins in this study, homeodomain proteins again clearly emerge as accurate descriptors of neuronal identity in the male-specific nervous system. The expression of homeobox genes also delineates neuronal subclasses within the previously known cardinal 23 male neuron classes. Such subclassification has been evident already based on terminal markers (e.g. among CA, CP or ray neurons), but for other classes, such subclassification was not previously appreciated (e.g. EF, DX neurons, class members distinguished by LIM-7 and UNC-4, respectively). We assigned at least one homeodomain protein to all 25 cardinal neuron classes of the male-specific nervous system of *C. elegans*, and, except for two closely related groups of neuron classes, we observed neuron specific combination of homeodomain proteins in each of these classes. The broad coverage of neuron classes with a limited number of examined homeodomain proteins predicts that a complete map of all homeodomain proteins will result in neuron type-specific combination of homeodomain proteins for each individual male-specific neuron class, as observed in the hermaphrodite nervous system.

As in the sex-shared nervous system, there is no obvious match of a specific neurotransmitter system with a specific homeobox signature, at least as far as the most commonly deployed neurotransmitter systems (ACh, Glu, GABA) are concerned, thereby corroborating the theme that neurotransmitter identity of distinct neuron classes is controlled in a “piece-meal” manner through distinct regulatory factors. This notion is further supported by the dissection of the *cis*-regulatory architecture of the *eat-4/VGluT* locus which identified distinct cis-regulatory elements driving gene expression in distinct male-specific neuron classes [62].

The HOX cluster homeodomain proteins, which constitute only 6 of the 102 homeodomain proteins in *C. elegans,* stand out in several regards. First, unlike in head ganglia, HOX gene expression very densely covers almost all neuron classes in male tail ganglia, oftentimes in an overlapping manner (summarized in **Fig 2**). This is a mirror image of many non-cluster homeobox genes that are expressed in multiple sex-shared head neuron classes but not at all in tail ganglia (**S1 Table**). Second, HOX cluster genes show clear anterior-posterior (a/p) patterns of collinearity within tail ganglia. Such co-linearity had already been observed in ventral cord neurons in *C. elegans* of sex-shared and sex-specific neurons [23, 63, 64]. Our systematic analysis of all HOX cluster genes extends such a/p-patterned expression throughout all neurons within male tail ganglia (i.e. the much-expanded lumbar ganglion in males, as well as the male-specific cloacal ganglion). This neuronal expression is – like all homeodomain protein expression patterns that we describe here – maintained throughout mature, adult stages.

### Homeobox gene function

Previous work has defined functions of HOX cluster genes in the male nervous system [16, 25, 29]. We have leveraged here the power of the NeuroPAL cell fate tool, in combination with additional cell fate markers that we developed, to describe the impact of removal of non-HOX cluster homeobox gene on the differentiation program of individual male-specific neurons. In two cases, notably PGA and MCM, we have shown that of the many neuron type-specific identity features that we tested, every single feature is affected in the respective homeobox gene mutant (*unc-30* for PGA; *unc-42* for MCM), arguing for a strict co-regulation of distinct identity markers, the key defining feature of a terminal selector-type regulatory logic [65]. In the case of MCM, we have also identified a homeobox gene, *ttx-1* which appears to act at an earlier step in the differentiation program by apparently priming the AMso glia cell to transdifferentiate into MCM.

We had previously noted that a subset of terminal selector demarcates neurons that are more highly interconnected with each other compared to other neurons and suggested that such terminal selectors may act as “circuit organizers” [49]. The best described of these cases is the *unc-42* homeobox gene which acts as a terminal selector in all 15 interconnected neurons in a nociceptive repulsive reflex circuit [49]. The function of *unc-42* as a terminal selector in the male-specific MCM neuron, the only sex-specific neuron in which *unc-42* is expressed in, fits into this theme since MCM is extensively connected to this *unc-42(+)* reflex circuit [9]. We therefore predict that in *unc-42* mutants, MCM may not only lose features of its molecular identity but may also fail to wire into the “*unc-42* circuit”. It is conceivable that UNC-42 may regulate the expression of cell surface molecules that allow for selective fasciculation of these neurons.

We have also described cases here in which it is either not clear whether a homeobox gene acts as a master-regulatory terminal selector or whether it rather regulates only individual aspects of a neuron function. In the DVE and DVF neurons, *lim-6* appears to control some, but clearly not all identity features. The effect of *lin-11* and *ttx-1* in the synaptically interconnected PVX, PVY and PHD neurons is also restricted to only a small number of identity features. However, these effects may have significant functional consequences, as evidenced by behavioral defects observed in *ttx-1* and *lin-11* mutants. *lin-11* and *ttx-1* may either act redundantly with other homeobox genes to specify PVX, PVY and PHD differentiation and/or other PVX/PVY/PHD-expressed factors may fulfill a master regulatory terminal selector role.

Akin to the mating defects that we describe here in *lin-11* and *ttx-1* mutants, mating defects have also been observed in another homeobox gene, *mec-3* [66, 67]. The sites of expression of *mec-3* that we described here match the function of the neurons and the specific behavioral defects in *mec-3* mutants (HOB neuron for vulva stopping; ray neurons for turning behavior)[67]. Yet, as in the case of *lin-11* and *ttx-1* mutants, cell type-specific rescue experiments are needed to confirm that these genes indeed act in these respective neurons to control animal behavior. Due to pleiotropies of other homeobox gene mutants (e.g. Unc phenotype of *unc-42* and *unc-30*, Let phenotype of *ceh-43,* Clr phenotype of *ceh-10* etc.), more cell type-specific knock-out approaches will be needed to assess their impact on the function in the male-specific neurons that these genes are expressed in.

### Limitations and conclusions

There are several limitations of the studies described here. The expression of half of all conserved homeodomain proteins still awaits investigation in the male-specific nervous system. Cell type-specific rescue experiments and/or cell type-specific knock-out experiments are currently lacking and are particularly needed to link the behavioral defects of *lin-11* and *ttx-1* mutants to their specific cellular focus of action. Such approaches will require the development of drivers with required cellular specificity. Also, while our cell fate marker analysis has clearly assigned function to several homeobox genes in identity specification, the analysis of more markers would be useful. Despite these limitations, we can conclude that homeobox gene expression and function appears to be as pervasive in the sex-specific nervous system as it is in the sex-shared nervous system.

These observations are in further support of the hypothesis that homeobox genes may have fulfilled an evolutionarily ancient function in neuronal differentiation [6].

## ACKNOWLEDGEMENTS

We thank Chi Chen for generating *C. elegans* mutant and transgenic strains and members of the Hobert lab for comments on this manuscript. G.V was supported by an EMBO Postdoctoral fellowship. This work was funded by the NIH (R01NS039996, R01NS137594) and the Howard Hughes Medical Institute.

## SUPPLEMENTARY FIGURES AND TABLES

**S1 Fig:**
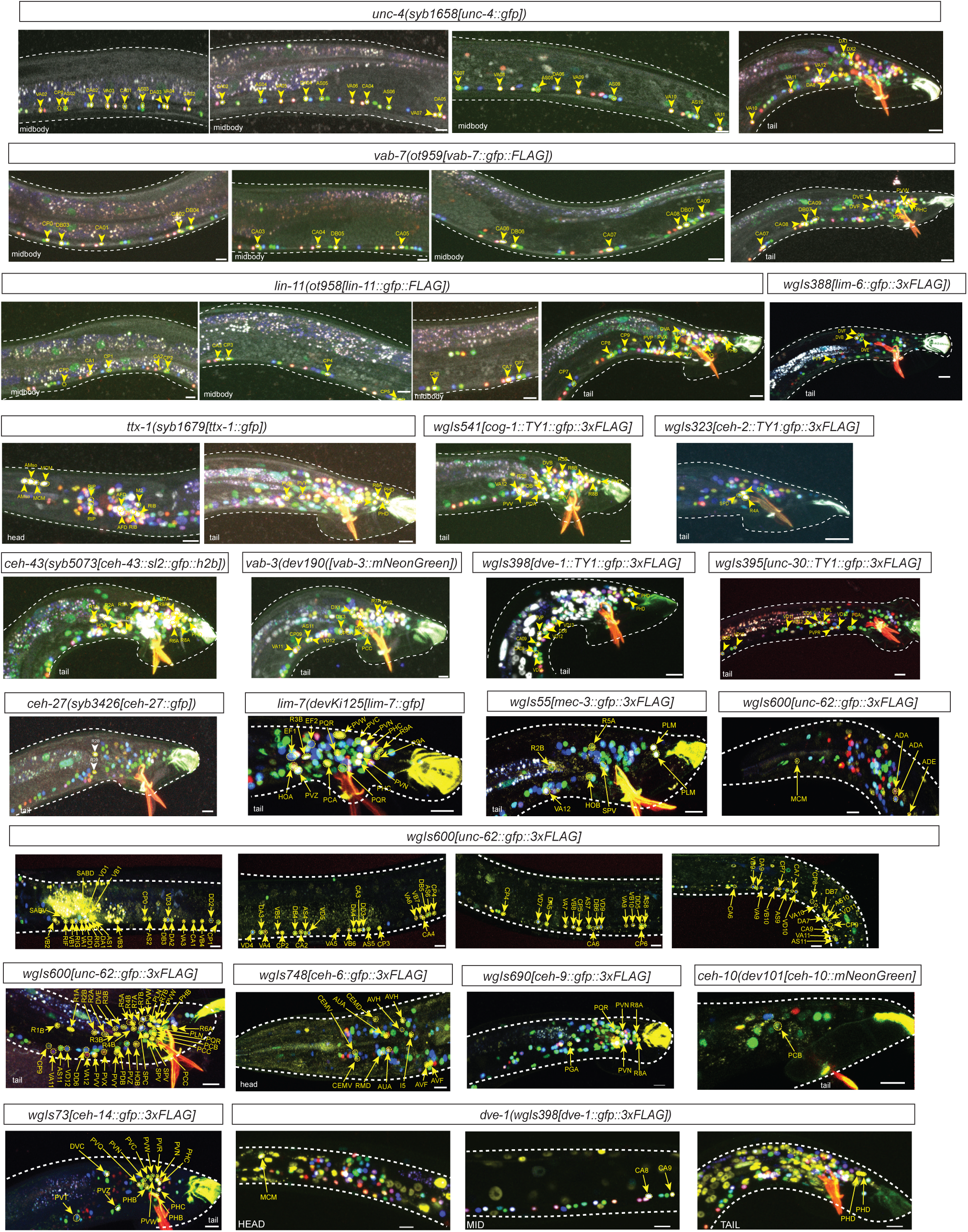
NeuroPAL ID data for homeobox reporter alleles. Homeobox reporter alleles from Fig 1 merged with NeuroPAL colors. Panels show representative images of NeuroPAL (*otIs669* or *otIs696*) landmark allele where all pseudocolors, as described [28, 29], are merged together with the GFP signal of homeobox reporters. Circles indicate the nuclear signal, whilst arrows are used to match the signal to the specific Neural ID. White dotted lines are used to trace the contours of the animal and its pharynx. Multiple images acquired with the NeuroPAL landmark are used to obtain a neuronal ID that is overlaid on representative images.

**S2 Fig:**
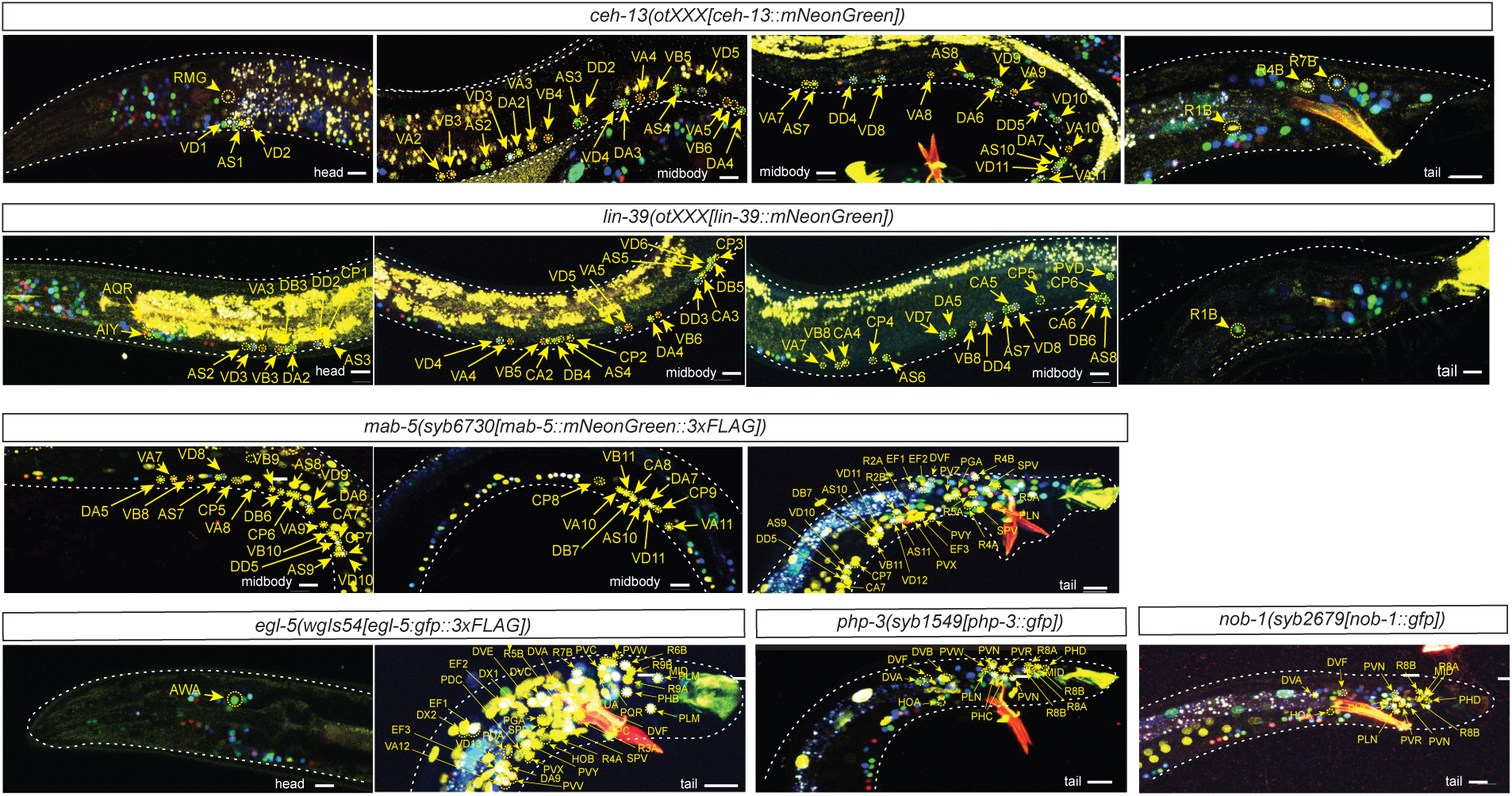
NeuroPAL ID data for HOX cluster alleles HOX cluster reporter alleles from Fig 2 merged with NeuroPAL colors. Panels show representative images of NeuroPAL (*otIs669* or *otIs696*) landmark allele where all pseudocolors, as described [28, 29], are merged together with the GFP signal of HOX reporters. Circles indicate the nuclear signal, whilst arrows are used to match the signal to the specific Neural ID. White dotted lines are used to trace the contours of the animal and its pharynx. Multiple images acquired with the NeuroPAL landmark are used to obtain a neuronal ID that is overlaid on representative images.

**S3 Fig:**
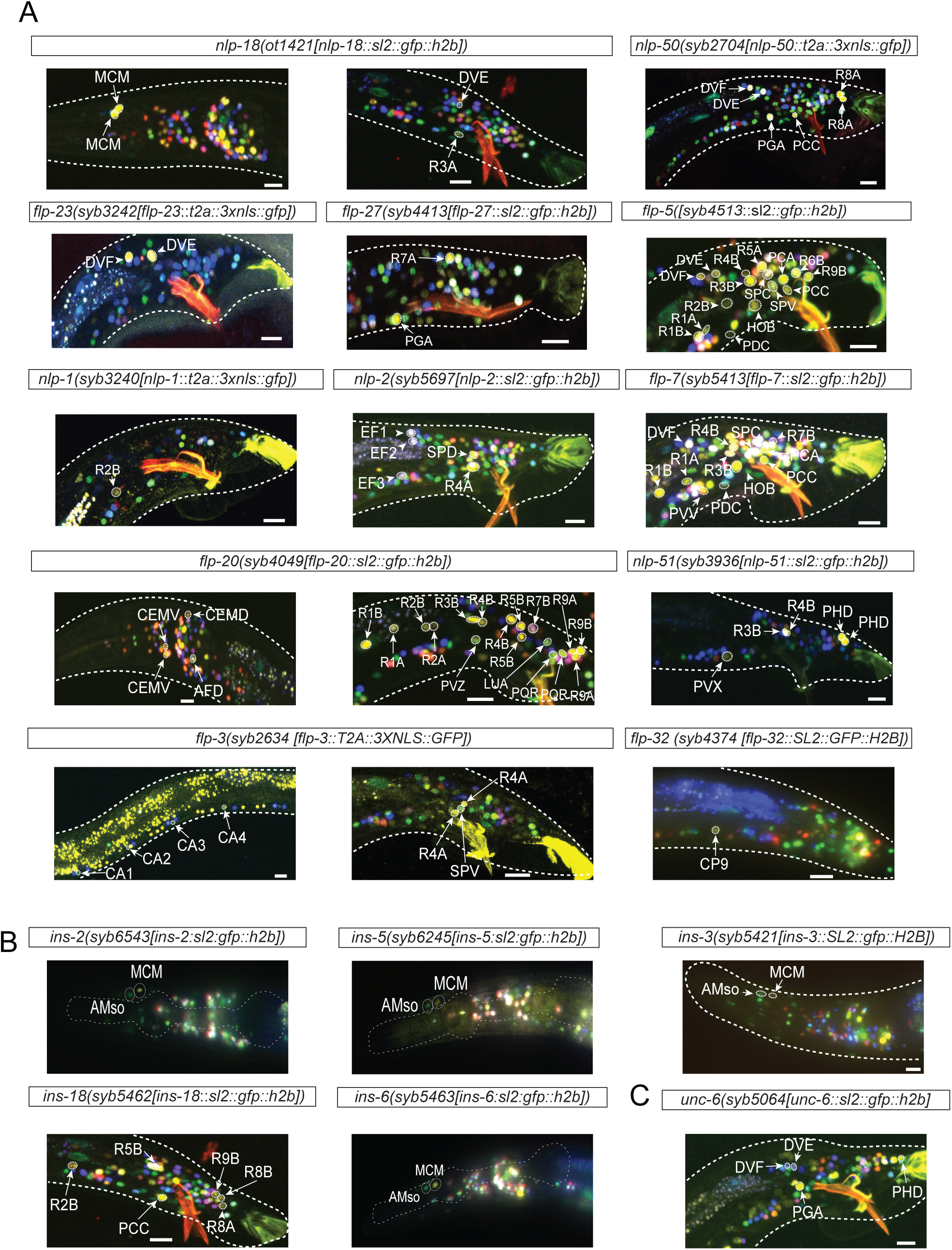
NeuroPAL ID data for new terminal identity markers for male-specific neurons. **A,B**: New terminal identity markers for male-specific neurons shown in Fig 4 merged with NeuroPAL colors. Panels show representative images of NeuroPAL (*otIs669* or *otIs696*) landmark allele where all pseudocolors, as described [28, 29], are merged together. Circles indicate the nuclear signal, whilst arrows are used to match the signal to the specific Neural ID. White dotted lines are used to trace the contours of the animal and its pharynx. Multiple images acquired with the NeuroPAL landmark are used to obtain a neuronal ID that is overlaid on representative images.

**S4 Fig:**
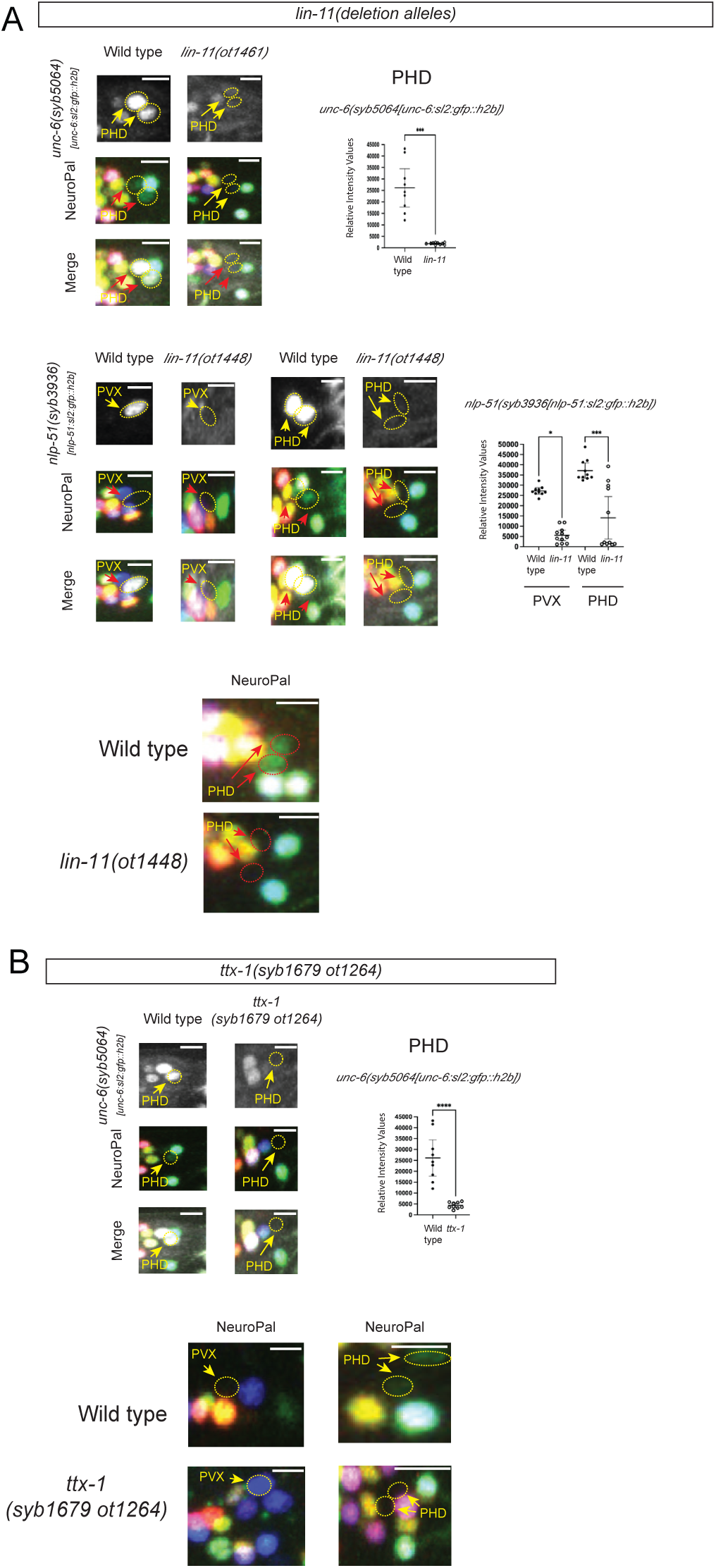
*ttx-1* and *lin-11* mutant analysis. **A:** *lin-11* full deletion mutants affect the expression *of unc-6/Netrin* in PHD. Representative images show the expression of *unc-6(syb5064)*, *nlp-51(syb3936)*, NeuroPAL colors(*otIs669*) and a merge of both in wild type and *lin-11(ot1448)* null mutants. Bar graph quantifies the decrease in Relative Intensity Value of *unc-6(*syb5064) and *nlp-51(syb3936)* expression in PHD and PVX, PHD respectively. **B:** *ttx-1* mutants affect the expression of *unc-6/Netrin* in PHD. Representative images show the expression of *unc-6(syb5064))*, NeuroPAL colors(*otIs669*) and a merge of both in wild type and *ttx-1(syb1679 ot1264)* mutants. Bar graph quantifies the decrease in Relative Intensity Value of *unc-6(syb5064)* expression in PHD. Statistics: *p value ≤0.05, ***p value ≤0.001, ****p value ≤0.0001, ns, not significant. Testing was performed using the Student t-test.

**S1 Table:**
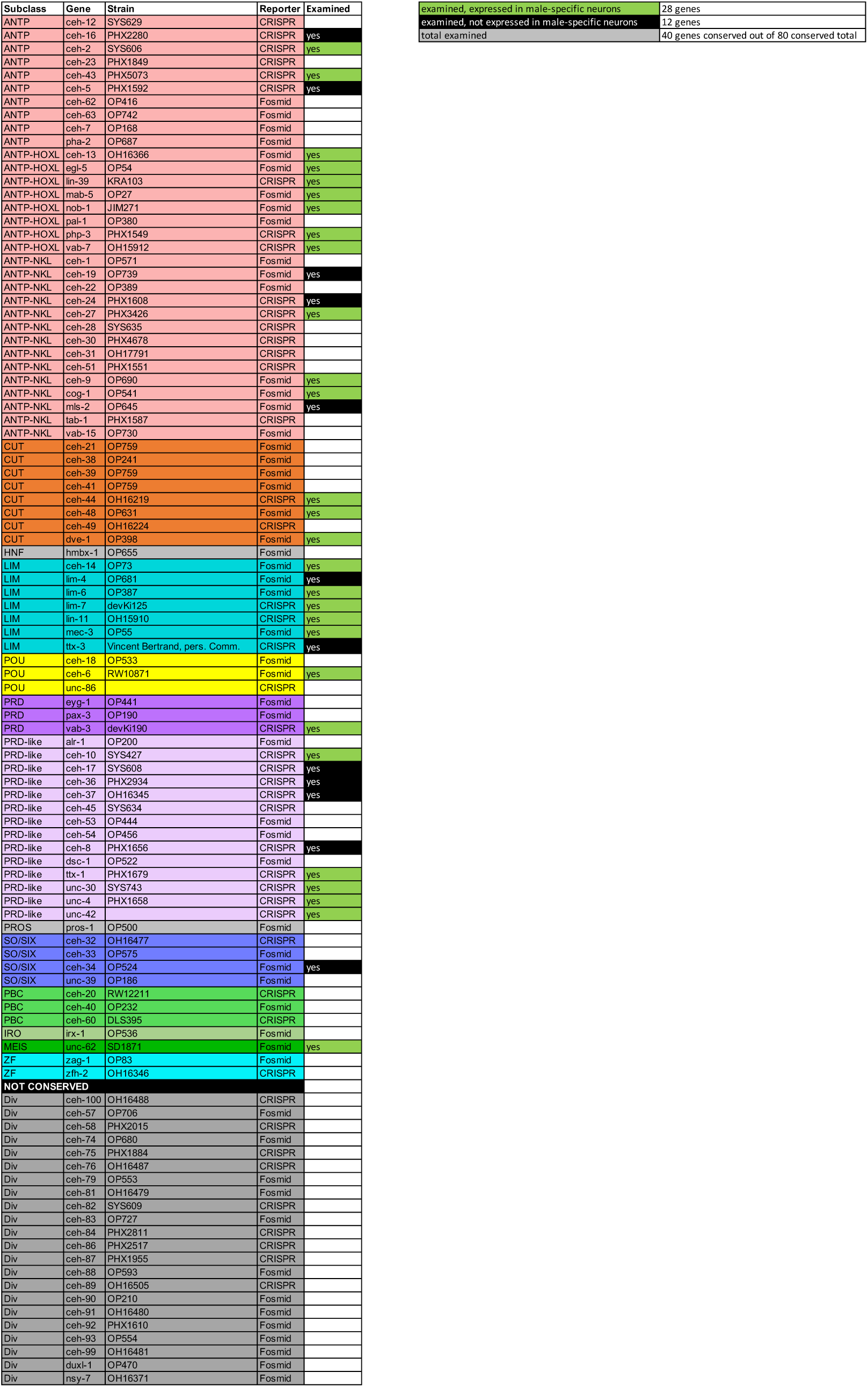
List of all *C. elegans* homeodomain proteins. The table indicates which of the 102 *C. elegans* homeodomain proteins were analyzed for expression in the male-specific nervous system.

**S2 Table:**
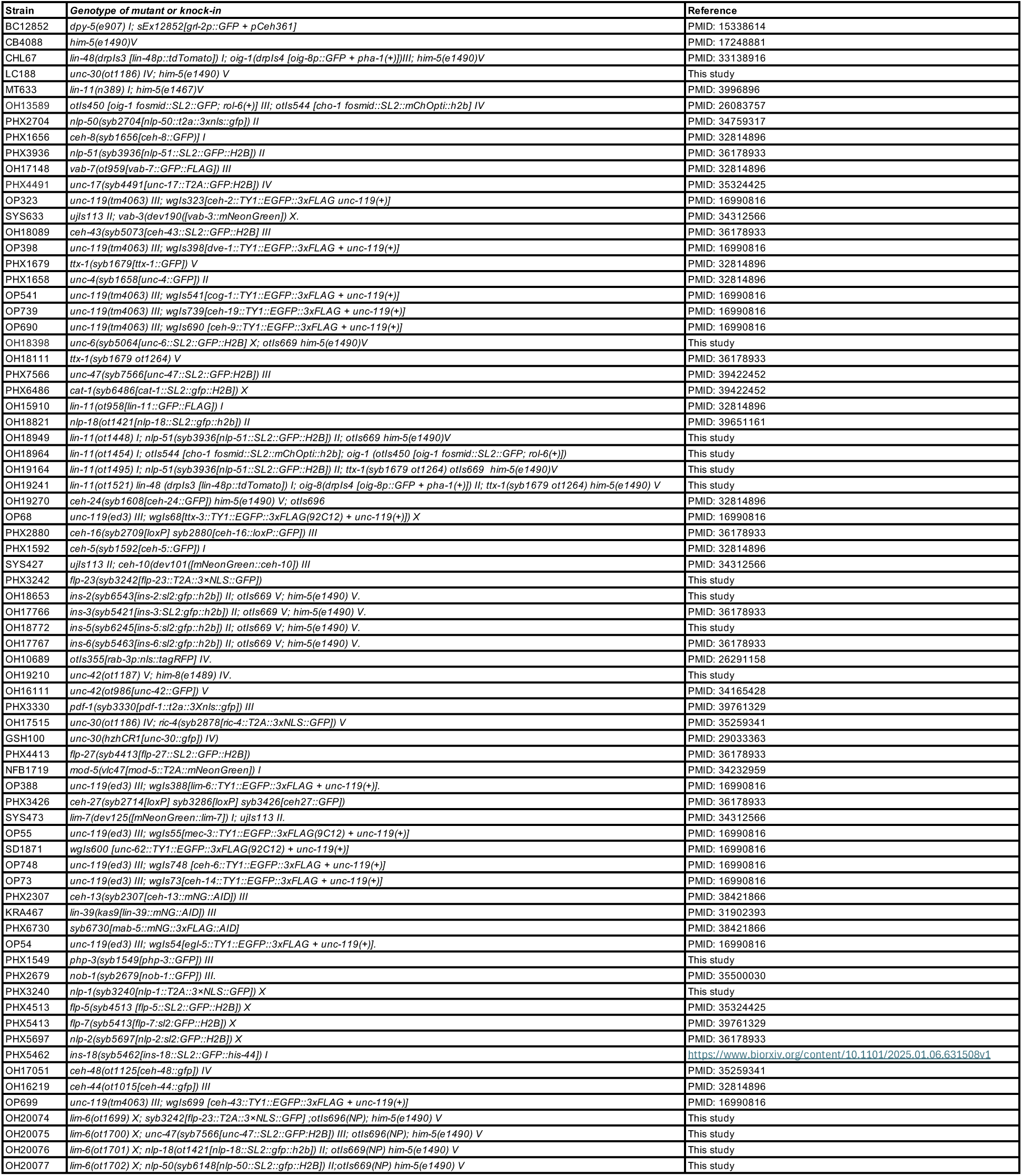
Strain list. List of all strains used in the manuscript.

